# Molecular Mechanisms of Cardiomyocyte Aging

**DOI:** 10.1101/2025.07.29.666777

**Authors:** Dogacan Yucel, Michael A. Trembley, Qingen Ke, Zexuan Wu, Peter Kang, William T. Pu

## Abstract

Aging is a major risk factor for cardiovascular diseases, yet the underlying molecular mechanisms remain poorly understood. In this study, we integrated physiological characterization of cardiomyocyte (CM) aging with concurrent single-nucleus RNA-seq and ATAC-seq, and reduced representation bisulfite sequencing to delineate the cellular and molecular landscape of CM aging in mice. Our analysis revealed significant age-associated changes in CM physiology, including hypertrophy, fibrosis, and diastolic dysfunction. We uncovered dramatic epigenetic remodeling in aged CMs, characterized by increased chromatin accessibility and altered DNA methylation patterns. Overexpression of the DNA methylase DNMT3A in young adult mouse hearts recapitulated key features of the aged heart phenotype, establishing DNA hypermethylation as a significant regulator of age-related CM function. Furthermore, ESRRG, an orphan nuclear receptor, functions as a mediator of diastolic function in the heart. Its overexpression significantly improved diastolic function and reduced expression of a non-coding RNA that is upregulated in aged CMs. These novel insights into the molecular mechanisms underlying cardiac aging identify molecular regulators involved in age-associated cardiac remodeling.

## Introduction

Advanced age is a major risk factor for the development of cardiovascular diseases (CVDs)^1^, including heart failure with preserved ejection fraction (HFpEF), which constitutes approximately half of all heart failure cases^1–3^. As the global population aged 50 and over is projected to increase by 61% between 2020 and 2050, and fertility rates remain at an all-time low, the prevalence of CVDs is projected to rise significantly^4^. This increase will impact individual health and pose a substantial economic burden on society^5^. To mitigate this impending crisis, it is crucial to understand the mechanisms of cardiac aging and why advanced age makes the heart more susceptible to CVDs.

Aging is characterized by well-established hallmarks that drive tissue dysfunction across multiple organs. These hallmarks include genomic instability, telomere attrition, epigenetic alterations, loss of proteostasis, impaired autophagy, cellular senescence, mitochondrial dysfunction, deregulated nutrient sensing, stem cell exhaustion, altered intercellular communication, and chronic inflammation^6,7^. Similarly, aging hearts exhibit distinct functional changes, most notably diastolic dysfunction^8,9^, characterized by impaired relaxation and increased stiffness of the left ventricle. While previous studies documented cellular changes in aged hearts, including cardiomyocyte (CM) hypertrophy, increased fibrosis, and altered mitochondrial function^10–12^, the molecular mechanisms driving these changes remain poorly understood. Particularly, the cell-type-specific alterations in gene expression and chromatin accessibility that occur during cardiac aging have not been comprehensively mapped. Key outstanding questions include: (1) how do transcriptional programs of different cardiac cell types respond to aging at the molecular level, (2) what are the transcriptional and epigenetic mechanisms that drive these changes in gene expression programs during cardiac aging, and (3) how do these changes increase susceptibility to heart disease.

To begin to gain insights into these questions, we performed concurrent single-nucleus RNA sequencing (snRNA-seq) and assay for transposase-accessible chromatin using sequencing (snATAC-seq) on young (4-months-old) and aged (28-months-old) cardiac tissues to comprehensively profile the transcriptomic and epigenomic landscapes of aging CMs. Through integrated analysis of these datasets, we discovered CM-specific molecular networks that are altered during aging. We identified and validated DNA methyltransferase 3a (DNMT3A) and estrogen related receptor gamma (ESRRG) as regulators of CM aging and age-related diastolic dysfunction.

## Results

### Phenotypic changes in CM aging

To identify CM-specific changes that occur with advancing age, hearts from mice at 4, 12, 18, 24, and 28 months were collected for physiological phenotyping (Fig. 1A). Consistent with previous studies, heart weight and heart weight to body weight (HW/BW) ratios significantly increased with age (Fig. 1B-C)^9,13^. Morphological analysis revealed an increase in CM cross-sectional area and a decrease in CM length/width ratio in older mice, indicating the development of age-associated concentric hypertrophy (Fig. 1D-E). This hypertrophic remodeling was further evidenced by an increase in anterior LV wall thickness in 28-month-old hearts compared to 4-month-old hearts (Supp. Fig. 1A).

**Figure 1.**
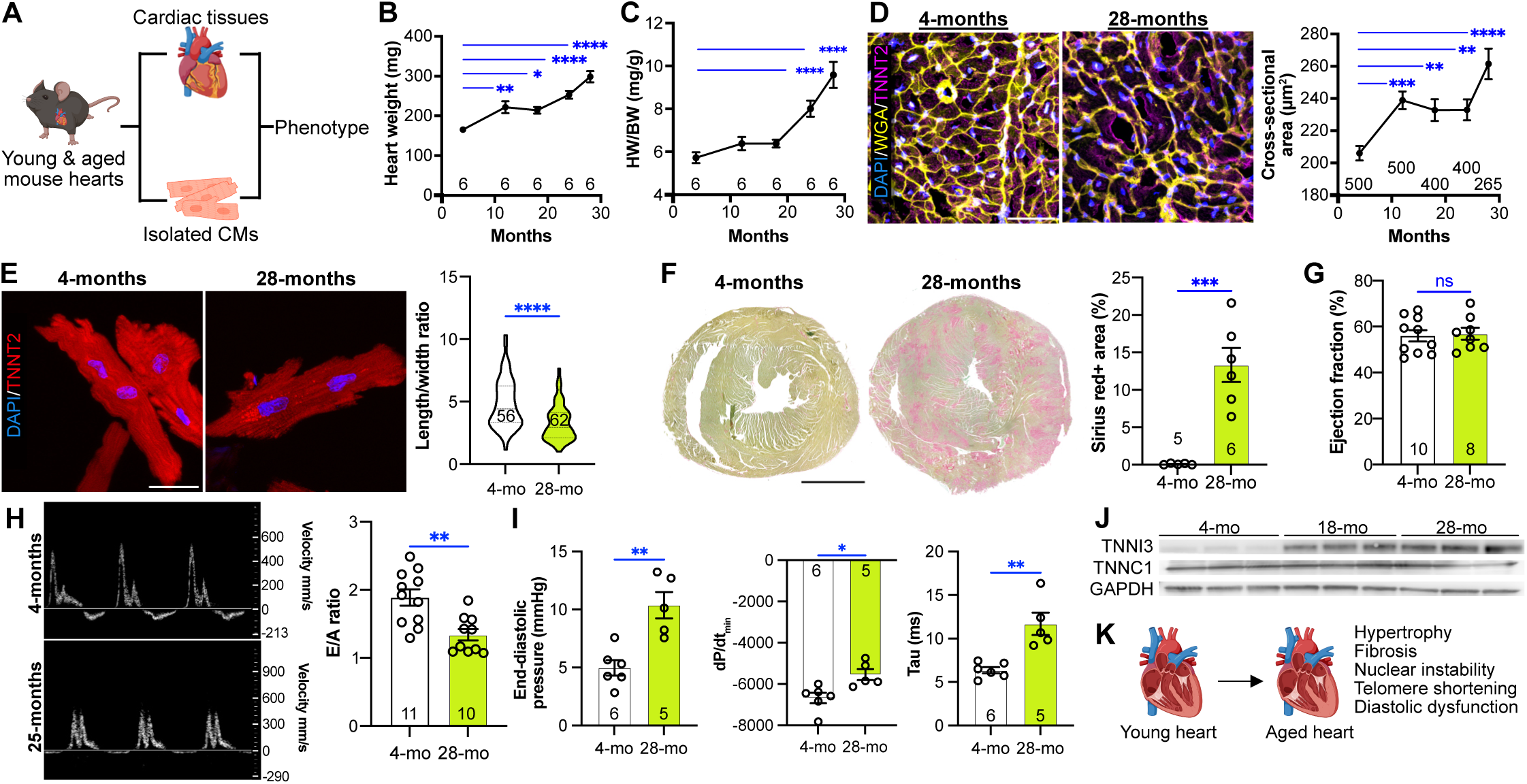
Age-associated changes in cardiac structure and function. **(A)** Summary of the experimental design. **(B-C)** Heart weight and heart weight normalized to body weight (HW/BW) across age groups (4, 12, 18, 24, and 28 months). One way ANOVA with Dunnett’s post hoc test. **(D)** Wheat germ agglutinin (WGA) staining of cardiac sections from 4- and 28-month-old mice. Left: Representative images. Bar, 50 μm. Right: Quantification of CM cross-sectional area. One way ANOVA with Dunnett’s post hoc test. **(E)** CM length/width ratio. Left: Representative images of isolated CMs. Bar, 25 μm. Right: Quantification. *t*-test. **(F)** Fibrosis assessment by picrosirius red/fast green staining. Left: Representative images. Bar, 1 mm. Right: Fibrotic area as a percentage of total tissue area. *t*-test. **(G-H)** Echocardiographic assessment of systolic (G, ejection fraction) and diastolic (H) heart function. Diastolic function was assessed using the ratio of early to late mitral inflow velocity (H, E/A ratio) in 4- and 28-month-old mice. *t*-test. **(I)** Invasive hemodynamic measurement of left ventricular diastolic function. Left: End-diastolic pressure. Middle: Minimum dP/dt. Right: time constant for isovolumic relaxation (tau). *t*-test. **(J)** Western blot analysis of TNNI3, TNNC1, and GAPDH protein levels in cardiac lysates from 4-, 18-, and 28-month-old mice. Representative of blots from 3 independent experiments. **(K)** Schematic summary of phenotypic changes observed in aged mouse hearts. Data are presented as mean ± SEM. Numbers within graphs indicate independent biological replicates. *p < 0.05, **p < 0.01, ***p < 0.001,****p<0.0001 ns = not significant. Panels A and K were created with the help of biorender.com.

Changes in nuclear morphology are frequently associated with aging and cellular senescence^14,15^. Immunostaining for PCM1, which labels the CM nuclear envelope, and DAPI demonstrated significant enlargement of CM nuclei in cardiac sections and isolated CMs from aged mice (Supp. Fig. 1B, D). In contrast, no significant changes in nuclear size were observed in the non-CM population (Supp. Fig. 1C). CM nuclei from aged mice also exhibited altered morphology, with increased roundness (Supp. Fig. 1E) and reduced solidity, a measure of the regularity of the nuclear boundary (Supp. Fig. 1F).

Histological assessment of fibrosis by picrosirius red and fast green staining revealed an increase in fibrotic area in hearts from 28-month-old compared to 4-month-old mice. Fibrosis occurred in a patchy rather than diffuse, interstitial pattern (Fig. 1F). This pattern often accompanies fibrotic replacement of dead CMs, suggesting increased cell death in aged hearts^16^. Immunostaining for γH2A.X also demonstrated increased prevalence of DNA damage foci in CMs from aged mice (Supp. Fig. 1G), consistent with greater DNA damage in aged CMs^17^.

While telomere shortening is well-documented in proliferating cells, its dynamics in post-mitotic cells like CMs are less understood^18,19^. Moreover, shortened telomeres have been associated with CM dysfunction, particularly under stress^20–22^. Relative telomere length, measured by qPCR, was lower in CM nuclei isolated from 28-month-old compared to 4-month-old hearts (Supp. Fig. 1H). Analysis of CM nuclear ploidy by flow cytometry showed no significant difference in the frequency of 2N vs 4N nuclei between young and aged hearts (Supp. Fig. 1I), suggesting that CM telomere shortening occurs independently of differences in cell cycle activity. At an earlier timepoint, a prior study demonstrated that length-independent telomere damage may play a significant role in cardiac aging and precede telomere shortening^23^. The telomere shortening that we observed at the more advanced stage of cardiac aging that we studied likely combines with accumulated telomere damage to contribute to advanced aging phenotypes.

Cardiac function in aged murine hearts is not well-defined, with some but not all studies showing altered systolic function^9,24–28^. Our echocardiographic assessment of cardiac function revealed that fractional shortening, a measure of systolic function, and heart rate did not differ significantly between 28-month-old and 4-month-old mice (Fig. 1G and Supp. Fig. 1J). Consistent with the gravimetric measurements, echocardiographic LV mass was greater in aged compared to young mice (Supp. Fig. 1K). Echocardiographic left ventricular wall thickness was significantly higher in the 28-month-old group (Supp. Fig. 1L). The ratio between early and late mitral inflow velocity (E/A ratio), a measure of diastolic function, was significantly lower in aged hearts (Fig. 1H), suggesting diastolic dysfunction, consistent with other reports^29–31^. Moreover, 28-month-old mice exhibited lower left ventricular end-diastolic volume (Supp. Fig. 1M), further suggesting impaired relaxation. We performed invasive intracardiac pressure-volume recordings, the gold standard for assessment of cardiac diastolic function, to confirm diastolic dysfunction in aged mice. End-diastolic pressure, dP/dt_min_, and the time constant for isovolumic relaxation (Tau) were all higher in aged hearts (Fig. 1I), confirming diastolic dysfunction.

Pathogenic variants in TNNI3, a component of the sarcomere thin filament, frequently cause restrictive cardiomyopathy, which is also characterized by diastolic dysfunction^32–34^. Western blot analysis demonstrated upregulation of TNNI3 with advancing age, while levels of TNNC1, another thin filament protein, were unchanged (Fig. 1J, Supp. Fig. 1N). Immunofluorescence microscopy revealed altered localization of TNNI3 in CMs from older mice, with increased perinuclear accumulation, suggestive of impaired sarcomere incorporation and/or protein aggregate formation (Supp. Fig. 1O). In contrast, immunostaining for titin, a critical determinant of myocardial passive stiffness, showed normal sarcomeric organization and localization in both isolated CMs and cardiac tissue sections from aged hearts (Supp. Fig. 1P-Q), suggesting that CM aging may involve altered localization, aggregation, or degradation of specific sarcomeric proteins rather than global sarcomeric disorganization.

In summary, our phenotypic characterization of hearts from mice at different ages revealed significant age-associated changes (Fig. 1K). Aged hearts were larger, more fibrotic, and had impaired diastolic function. CMs displayed concentric hypertrophy with nuclear enlargement and telomere shortening. These physiological and cellular phenotypes provide a solid foundation for investigating the molecular mechanisms underlying cardiac aging.

### Molecular changes in aged ventricles

To correlate functional and morphological alterations in aged cardiac cells with molecular signatures, we assessed the RNA expression of established general aging markers associated with senescence-associated secretory phenotype (SASP), DNA damage repair, cellular senescence, telomere maintenance, metabolism, autophagy, stress response, and inflammation in young and aged cardiac ventricles (Supp. Fig. 2). Contrary to expectations, the majority of general aging markers did not significantly differ between age groups. The exceptions were *Sod1*, *Sod2*, *Cxcl1*, *Parp1*, and *Sirt1*. The downregulation of *Sod1* and *Sod2* suggests dysregulation of oxidative stress management in aged ventricles. Interestingly, we did not detect statistically significant differences in the expression of canonical senescence markers such as p16 and p21, indicating that these established general aging biomarkers do not reliably reflect cardiac aging, underscoring the need for tissue-specific markers of cellular aging.

### Single-nucleus transcriptomic and epigenomic profiling of cardiac aging

To investigate the molecular pathways of age-associated cardiac remodeling, we performed concurrent snRNA-seq and snATAC-seq on nuclei isolated from young (4 mo) and aged (28 mo) hearts in biological duplicate (Fig. 2A). By providing complementary information on gene expression and chromatin accessibility, respectively, this approach yielded an unprecedented view of the transcriptomic and epigenomic changes that occur during cardiac aging. After stringent quality control, we obtained 28,671 high-quality nuclei (young:13,814; old: 14,857). Quality control analyses demonstrated good coverage in ATAC and RNA values, robust TSS enrichment with clear nucleosome signals, and a strong correlation between TSS enrichment and ATAC counts (Supp. Fig. 3A-C), validating the quality of our snATAC-seq data. Integrated RNA and ATAC profiles were clustered and then visualized using Uniform Manifold Approximation and Projection (UMAP) (Fig. 2B). Clusters corresponded to major cardiac cell types, including CMs, endothelial cells (ECs), fibroblasts (FBs), pericytes (PCs), and immune cells, based on marker gene expression (Fig. 2C). Importantly, each replicate within the young and aged groups showed high consistency in both RNA and ATAC profiles, demonstrating the high quality and reproducibility of our data (Supp. Fig. 3D-E). Young and aged hearts segregated within cell type clusters, indicating distinct age-related transcriptomic and epigenomic signatures (Supp. Fig. 3F). The number of RNA features and counts in each cluster was similar between age groups (Supp. Fig. 3G-H), whereas there were more ATAC features and counts in aged nuclei for CMs, ECs, and FBs, and less for B-cells and T-cells (Suppl. Fig. 3I-J). Increased chromatin accessibility has been associated with epigenetic erosion and loss of cell identity during aging^35,36^.

**Figure 2.**
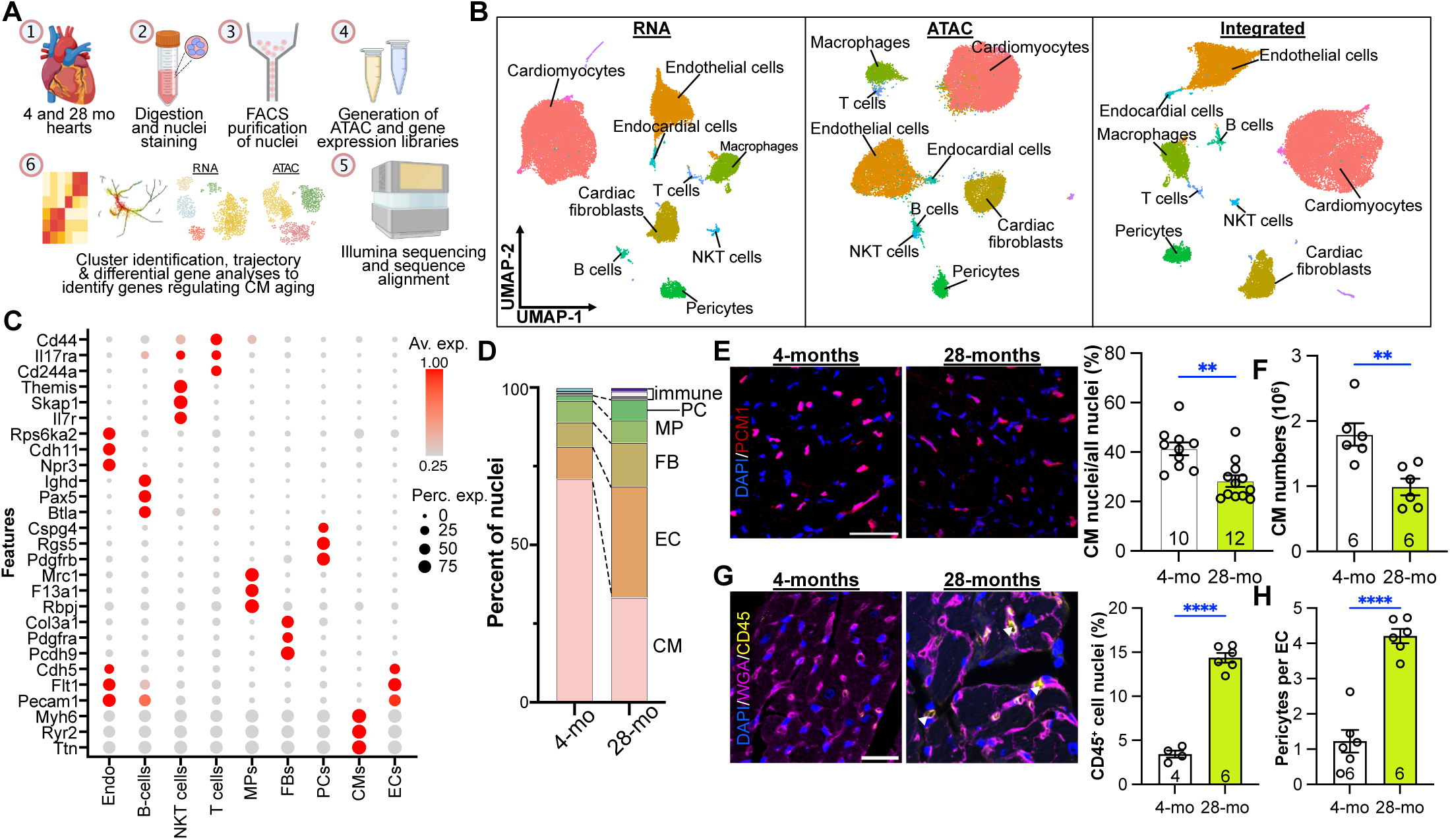
Single-nucleus transcriptomic and epigenomic profiling of cardiac aging. **(A)** Representative workflow of the single-nucleus experiment. Created with the help of biorender.com. **(B)** UMAP plots showing clustering of cardiac cell populations based on snRNA-seq data (left), snATAC-seq data (middle), and integrated analysis (right). **(C)** Expression of marker genes across different cell clusters. **(D)** Relative frequencies of cell types in young and aged cardiac populations from multiome analysis. **(E)** Cardiomyocyte (CM) nuclei analysis. Left: Representative images of PCM-1 immunostaining in cardiac sections from young and aged hearts. Scale bar, 50 μm. Right: Quantification of PCM-1+ CM nuclei (n=5 for 4-mo, n=6 for 28-mo groups). *t*-test. **(F)** Quantification of total CM numbers in young and aged hearts using isolated CM counting method (n = 6 mice per group). *t*-test. **(G)** CD45+ leukocyte analysis. Left: Representative immunohistochemical staining for CD45 in young and aged cardiac sections. Scale bar, 100 μm. Right: Frequency of CD45+ cell nuclei (%) (n=4 for 4-mo, n=6 for 28-mo groups). *t*-test. **(H)** Quantification of the ratio of pericytes to endothelial cells in young and aged hearts (n=6 mice per group). *t*-test. Data are presented as mean ± SEM. **p < 0.01, ****p<0.0001.

Quantification of the distribution of nuclei among cell types between young and aged hearts revealed that the proportion of CMs decreased and FBs increased in aged hearts (Fig. 2D). This age-associated reduction in the proportion of CM nuclei was validated by immunostaining for PCM1 (Fig. 2E) and flow cytometry of isolated CM nuclei (Supp. Fig. 3K). Furthermore, CMs isolated from fixed hearts, a method that minimizes CM loss during isolation^37^, confirmed the age-related decline in total CM numbers (Fig. 2F). The age-associated loss of CMs likely contributes to remodeling and increased disease susceptibility of the aged heart.

Consistent with the snRNAseq analysis, immunohistochemical staining revealed a higher number of CD45+ leukocytes in aged hearts compared to young hearts (Fig. 2G). Inflammation is an established hallmark of aging^6^, and it disrupts the homeostasis of several tissues^38^. While snRNA-seq suggested an expansion in the proportion of both ECs and PCs in aged hearts, immunostaining using isolectin B4 and NG2, markers of ECs and PCs, respectively, revealed that the number of ECs per CM was comparable between young and aged hearts (Supp. Fig. 3L), whereas the ratio of PCs to ECs was significantly higher in aged hearts, indicative of relative PC expansion during aging (Fig. 2H). This apparent discrepancy between single-nucleus sequencing and histological quantification likely arises from technical limitations inherent to snRNA-seq, such as differential efficiency of nuclear isolation across cell types and ages.

In summary, we observed altered cellular composition in aged hearts, characterized by reduced proportion of CMs and increased proportions of FBs, PCs, and CD45+ immune cells. These findings highlight the complex interplay between different cell types during cardiac aging and provide valuable insights into potential cellular mechanisms underlying age-related cardiac dysfunction.

### Cellular interplay in young and aged hearts

To examine how aging affects intercellular communication in the heart, we analyzed putative intercellular signaling pathways between major cardiac cell populations in young and aged hearts using the CellChat algorithm^39^. Our analysis revealed that the total number and strength of intercellular interactions were lower in aged cardiac cell populations (Supp. Fig. 4A), suggesting that transcriptional changes during aging compromise direct cell-cell communication networks. When examining specific intercellular interactions, we observed that the majority of cardiac cell populations in aged hearts exhibited reduced capacity to send signals to other cells, while certain immune cell types, particularly macrophages (MPs) and natural killer T (NKT) cells, showed enhanced capacity to receive signals from other cell types (Supp. Fig. 4B-C).

Analysis of specific ligand-receptor pairs revealed distinct age-associated changes in intercellular communication (Supp. Fig. 4D). Notably, ligand-receptor interactions involved in leukocyte proliferation were significantly enriched in aged cardiac tissue. Given these pronounced changes in immune cell interactions, we further characterized the cell-cell communication networks involving different immune cell populations. Young NKT cells primarily communicated through ECM-associated ligands (Collagen, Laminin) and growth factors (VEGF, IGF, NRG), whereas aged NKT cells engaged in interactions involving inflammatory proteins (SIRP, SELPLG, PECAM1, CD86, SELL) (Supp. Fig. 5A). This shift suggests a transition from structural support to increased inflammatory signaling with cardiac aging.

Macrophages, which constitute the most abundant immune cell population in both young and aged hearts, exhibited significant upregulation of extracellular matrix (ECM)-related ligand-receptor interactions with other cell types with age (Supp. Fig. 5B). Interestingly, both B cells and T cells engaged in more robust intercellular communication in young hearts compared to aged hearts, with the notable exception of the SIRP ligand-receptor axis in T cells, which has been previously implicated in regulating T cell activation (Supp. Fig. 5C-D).

To identify specific differential cell-cell interactions, we performed differential interaction analysis (Supp. Fig. 5E). The most striking change was observed in the ligand-receptor-mediated communication between cardiac fibroblasts (as signal-sending cells) and macrophages (as signal-receiving cells), with significantly stronger interaction in aged hearts compared to young hearts. Further analysis of this specific cell-cell interaction revealed enhanced Collagen and Laminin signaling by fibroblasts being received by macrophages via the Itga9 and Itgb1 receptors in aged hearts (Supp. Fig. 5F). Notably, other intercellular connections involving these cell types did not show such pronounced differences, highlighting the specificity of the fibroblast-to-macrophage communication axis in cardiac aging.

Finally, we examined intercellular interactions directly involving CMs in young versus aged hearts. While the majority of ligand-receptor interactions incoming to CMs remained relatively unchanged with age, we observed substantial alterations in direct cell-cell communication from CMs, B cells, and T cells to CMs (Supp. Fig. 6). These age-associated changes in intercellular interaction networks may contribute to the functional and structural remodeling observed in aged hearts and provide potential targets for interventions aimed at mitigating age-related cardiac dysfunction.

### Transcriptional signatures of aged CMs

Subclustering of CM nuclei revealed 6 transcriptionally distinct subpopulations (CM0-CM5; Fig. 3A). These CM clusters were differentially represented between young and aged hearts, with CM0 and CM1 constituting the majority of young and aged CM nuclei, respectively (Fig. 3B). Notably, each replicate within the young and aged groups showed high consistency in both RNA and ATAC profiles, demonstrating the reproducibility of our findings (Supp. Fig. 7A-B).

**Figure 3.**
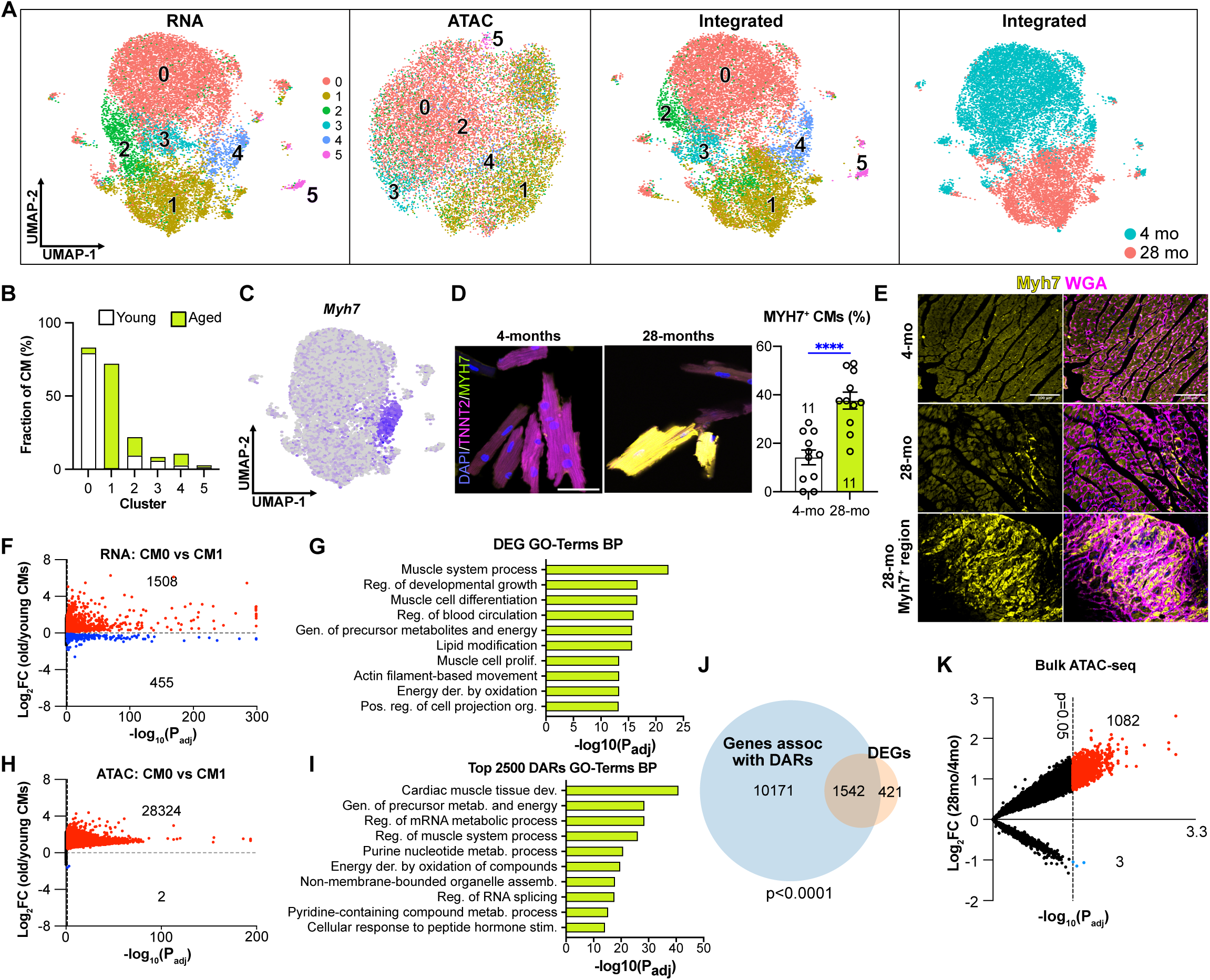
Transcriptional and epigenomic signatures of aged vs young cardiomyocytes. **(A)** UMAP plots of cardiomyocytes (CMs). From left to right: RNA, ATAC, and integrated UMAP plots, and the separation of 4- and 28-month CMs in the integrated UMAP. **(B)** Frequency distribution of CM clusters in young and aged hearts. **(C)** *Myh7* expression in CMs, highlighting cluster 4 as the *Myh7*+ population. **(D)** MYH7 immunostaining of isolated CMs. Left: Representative images. Scale bar, 50 μm. Right: Quantification. *t*-test. **(E)** MYH7 and WGA co-staining of cardiac sections, showing MYH7+ cells primarily in focal regions of 28-month-old hearts. Scale bar, 100 μm. **(F)** CM0 vs CM1 differentially expressed genes (DEGs). **(G)** Biological Process (BP) GO term analysis for CM0 vs CM1 DEGs. **(H)** CM0 vs CM1 differentially accessible regions (DARs). **(I)** GO term analysis of biological processes enriched within the top 2,500 DARs. **(J)** Overlap between DEGs and genes associated with DARs. Hypergeometric test. **(K)** Bulk ATAC-seq data comparing young and aged hearts. ****p < 0.0001.

Previous studies reported increased expression of cardiac stress markers, such as *Myh7*, *Nppa*, *Nppb*, and *Ankrd1*, in aged hearts^27,40^. RTqPCR on bulk RNA samples from young and aged cardiac tissues confirmed significant upregulation of these markers during aging (Supp. Fig. 7C). To determine if this upregulation occurs uniformly across all CMs or in a specific subset, we examined the expression of these markers in the CM subclusters. CM4 exhibited high levels of *Myh7, Nppa, Nppb, and Ankrd1* expression and was significantly expanded in aged hearts (Fig. 3B-C and Supp. Fig. 7C-D). Immunostaining of isolated CMs confirmed the presence of a MYH7-positive CM subpopulation (Fig. 3D), which was preferentially localized to fibrotic regions in aged hearts (Fig. 3E). Gene ontology analysis revealed that this "remodeled" CM cluster was associated with pathways related to cardiac contraction and development (Supp. Fig. 7E).

To identify age-related transcriptional changes, we compared the gene expression profiles of the predominantly young and old CM clusters, CM0 and CM1, respectively. Differential expression analysis revealed a higher number of upregulated compared to downregulated genes in aged CMs (Fig. 3F and Table S1). Biological process gene ontology terms enriched for CM0 vs. CM1 differentially expressed genes (DEGs) included those related to metabolism, contraction, and actomyosin organization (Fig. 3G). Notably, 9 out of the top 11 DEGs upregulated in aged hearts encoded mitochondrial proteins (Table S1). Aged CMs exhibited increased mitochondrial DNA content (Supp. Fig. 7F) and abnormal clustering of mitochondria (Supp. Fig. 7G), underscoring the prominent role of mitochondria in CM aging, similar to previous studies^41^. However, young and aged CMs had comparable mtDNA mutation frequency (Supp. Fig. 7H). Together these observations suggest that changes in mitochondria play a prominent role in CM aging.

Molecular function gene ontology terms enriched for CM0 vs. CM1 DEGs related to acyl-CoA activity, electron transfer chain activity, actin binding, cAMP binding, estrogen response element binding, and serine/threonine kinase activity (Supp. Fig. 7I). Some of these pathways complement the biological process changes, such as alpha-actinin and actin binding, which are connected to the heart contraction biological process term. Additionally, changes in the acetyl-CoA and PDE pathways provide more specific insights into which aspects of metabolism are affected.

### Epigenomic remodeling of the aged CM

snATAC-seq profiling revealed dramatic remodeling of the chromatin landscape of aged CMs. We found that 28,324 differentially accessible regions (DARs) had significantly greater accessibility, and only 2 DARs had lower accessibility in aged compared to young CMs (Fig. 3H-I and Table S2). DARs were highly enriched within promoter regions (Supp. Fig. 8A). Approximately 75% of differentially expressed genes (DEGs) overlapped DAR-associated genes (Fig. 3J), and chromatin accessibility at promoter regions positively correlated with gene expression changes during CM aging (Supp. Fig. 8B). This increased chromatin accessibility in aged CMs has been linked to dysregulation of the transcriptional machinery, epigenetic erosion, and partial loss of cell identity in aged CMs in other aging cell types^42,43^.

Similar to DEG GO terms, analysis of the top 2,500 DARs identified enrichment of biological process functional terms related to RNA transcription, cardiac contraction, oxidation and metabolism (Fig. 3I). Dysregulation of the transcriptional machinery and altered expression of genes involved in the metabolism, previously associated with aging in other cell types^44–47^, suggest that these pathways and those related to heart contraction contribute to cardiac aging. Similarly, analysis of the top 2,500 DARs identified enrichment of molecular function terms such as RNA/DNA binding, transcription factor binding, electron transfer activity,and serine/threonine kinase activity (Supp. Fig. 8C). Transcription factor binding, magnesium binding, electron transfer activity, and serine/threonine kinase activity were also identified in the DEG GO term analysis, reinforcing their potential importance in cardiac aging. Motif analysis identified several CpG-rich transcription factor binding motifs enriched within the DARs (Supp. Fig. 8D). Among these was the motif of Nrf1, which has been implicated in regulating CM redox balance and proteostasis.^48^

To further confirm these extensive changes in chromatin accessibility, we performed bulk ATAC sequencing on nuclei isolated from young and aged ventricles. Quality control metrics for these bulk ATAC-seq samples were comparable between groups (Supp. Fig. 8E). Consistent with the snATAC-seq results, bulk ATAC-seq demonstrated a significant global increase in chromatin accessibility in aged compared to young hearts, with 1082 DARs with greater and only 3 with lower accessibility in aged hearts (Fig. 3K). Five different normalization methods yielded a similar result (Supp. Fig. 8F), indicating that the striking increase in accessibility in aged hearts was not caused by biases introduced by different normalization approaches.

Another epigenetic mark linked to aging is DNA methylation at cytosine residues, particularly within CpG islands. DNA methylation is a well-established biomarker of aging, with biological clocks based on methylation patterns correlating strongly with chronological age^49,50^. Interestingly, DARs between young and aged CMs were significantly enriched for CpG islands (Fig. 4A). To further examine the changes in DNA methylation between young and aged CM nuclei, we performed reduced representation bisulfite sequencing (RRBS; Fig. 4B). Aged CMs exhibited increased DNA methylation in CpG (Fig. 4C) and CHG, but not CHH, contexts (Supp. Fig. 8G). 1893 differentially methylated regions (DMRs) had greater methylation in aged CMs compared to young CMs, compared to only 238 with less methylation (Fig. 4D and Table S3).

**Figure 4.**
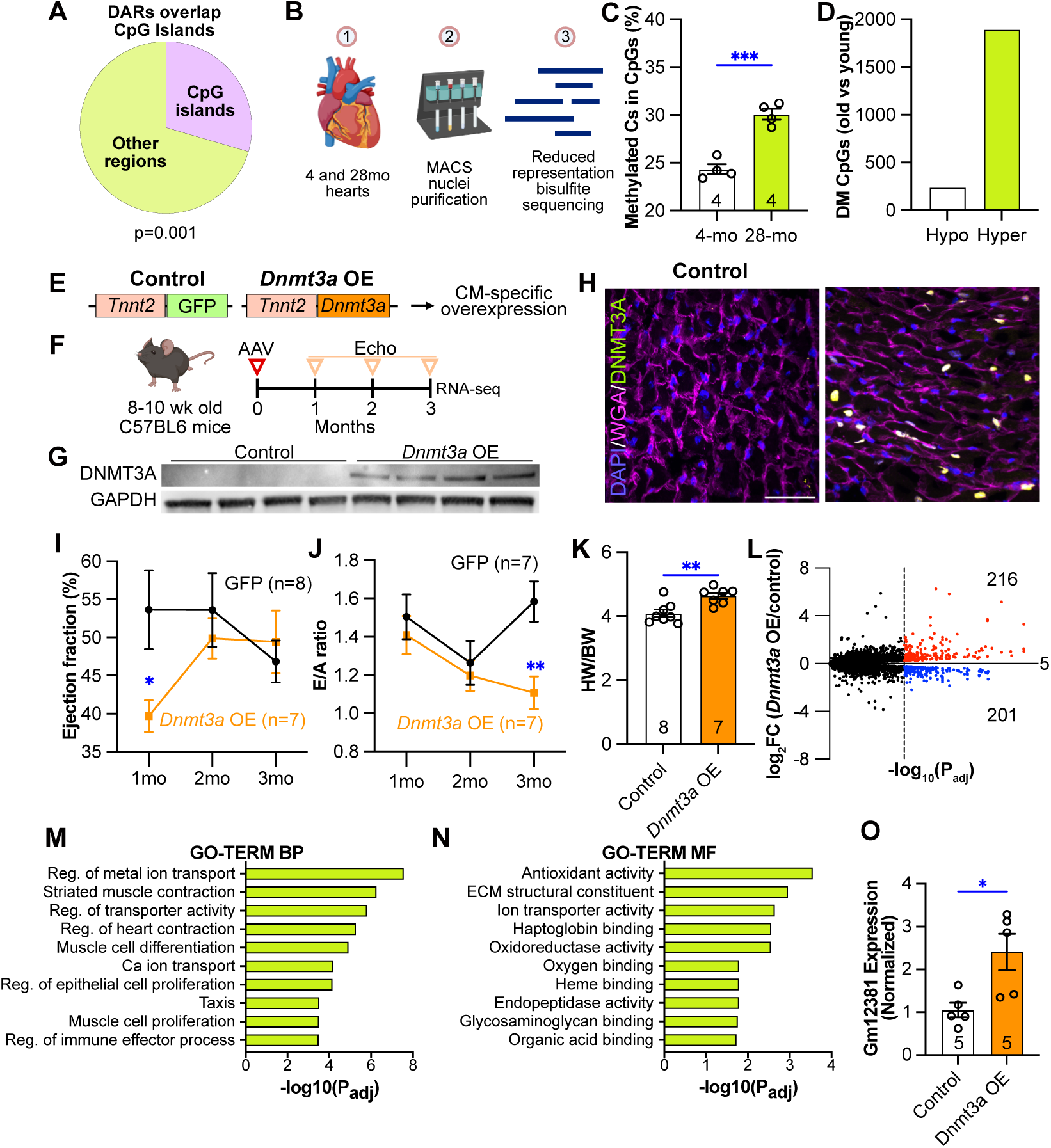
Epigenomic remodeling and functional consequences of DNA hypermethylation in aged cardiomyocytes. **(A)** ∼30% of DARs were in CpG islands. Permutation test. Data are presented as mean ± SEM where applicable. **(B)** Schematic representation of the reduced representation bisulfite sequencing (RRBS) experiment. **(C)** Graph showing percentage of methylated cytosines in CpG contexts for young and aged hearts. **(D)** Counts of hypomethylated and hypermethylated regions between 4- and 28-month-old hearts. **(E)** Schematic representation of AAV constructs for *Dnmt3a* overexpression (OE). **(F)** Timeline of AAV injection and experimental procedures. **(G)** Western blot analysis of DNMT3A protein levels in *Dnmt3a*-overexpressing (OE) and control hearts. **(H)** Immunohistochemical staining for DNMT3A in *Dnmt3a* OE and control heart sections. Scale bar, 50 μm. **(I)** Ejection fraction of *Dnmt3a* OE and control mice over 3 months. t-test. **(J)** E/A ratio measurements of *Dnmt3a* OE and control mice over 3 months. t-test. **(K)** Heart weight to body weight (HW/BW) ratios of *Dnmt3a* OE and control mice after 3 months. t-test. **(L)** Differentially expressed genes (DEGs; p value<0.05) between *Dnmt3a* OE and control hearts. **(M-N)** Gene ontology biological process (M) and molecular function (N) terms with genes over-represented among DEGs. (**O**) Expression level of *Gm12381* in *Dnmt3a* OE and control hearts 3 months after injection. Data are presented as mean ± SEM. t-test. *p < 0.05, **p < 0.01, ***p < 0.001. Panels B and F were created with the help of biorender.com.

The DMRs were enriched in biological processes related to TGF-beta, WNT and BMP signaling pathways, metabolism, actin structure, and cell-cell junction organization (Supp. Fig. 8H). Interestingly, the biological process GO terms associated with DMRs showed limited overlap with those from DARs and DEGs, with the notable exception of metabolic processes, which were consistently enriched across all datasets, highlighting the central role of metabolism in cardiomyocyte aging. The DMRs were enriched in molecular function terms related to DNA binding (Supp. Fig. 8I), which was also present in the DAR and DEG GO terms.

In summary, our multi-omics approach revealed extensive epigenomic remodeling in aged CMs, characterized by increased chromatin accessibility and altered DNA methylation patterns. These epigenomic changes were associated with widespread gene expression changes and may contribute to the age-related decline in CM function.

### Molecular changes in endothelial cells and fibroblasts

We analyzed the RNA and ATAC snRNAseq data to determine the molecular changes that occur in ECs and FBs during aging. FBs showed 263 DEGs (168 up and 95 down) between aged and young samples (Supp. Fig. 9A). Strikingly, we also identified more than 1600 regions that were more accessible in aged FBs compared to young FBs (Supp. Fig. 9B). In comparison, in ECs most genes were downregulated with age (154 down and 80 up; Supp. Fig. 9C). However, like CMs and FBs, ECs exhibited increased chromatin accessibility with age (Supp. Fig. 9D). CFBs exhibited broad alterations in extracellular matrix organization, cytoskeletal dynamics, and cell-matrix interactions, consistent with their structural and remodeling roles (Supp. Fig. 9E). In contrast, ECs showed enrichment in immune-related functions, protein transport, and signaling pathways including antigen presentation and cell adhesion (Supp. Fig. 9F). Despite these differences, shared pathways such as cAMP signaling, and metabolic activity were evident in both cell types, suggesting common regulatory mechanisms that may contribute to the aging or remodeling process across cell populations.

### Long non-coding RNAs as molecular markers of cardiac aging

A major role is emerging for lncRNAs in aging and senescence^51^. Using our multiomics data, we identified two lncRNAs, *Gm47283* and *AC149090.1*, that were consistently upregulated in aged cells of all types (Supp. Fig. 10A). To identify CM-specific aging biomarkers, we focused on lncRNAs differentially expressed between young and aged nuclei from CMs but not other cell types (Supp. Fig. 10B). Among these, *Gm12381* exhibited progressive upregulation in the heart with age (Supp. Fig. 10C). Importantly, *Gm12381* was induced in an age-dependent manner but not in response to stress induced by chronic isoproterenol infusion, suggesting its potential utility as a specific biomarker of CM aging (Supp. Fig. 10D-E).

### Causal link between DNA hypermethylation and age-related CM dysfunction

To understand how DNA methylation, chromatin accessibility, and gene expression in CMs correlate during aging, we integrated our RRBS, snATAC-seq, and snRNA-seq datasets (Table S4). For consistent comparison with our RRBS data, we performed pseudobulk analysis to compare all young versus aged CM nuclei, rather than the cluster-specific analysis shown in Figure 3. There were 558 genes that neighbored either DARs or DMRs, while 468 DEGs neighbored only DARs. Only 92 DEGs neighbored both DARs and DMRs (Supp. Fig. 11A).

We analyzed the correlation between aging-associated changes in accessibility, methylation, and gene expression. DARs showed a significant positive correlation with neighboring DEGs, suggesting that increased chromatin accessibility directly influences gene expression (Supp. Fig. 8B). We observed a significant negative correlation between DMRs and their neighboring DEGs (Supp. Fig. 11B), suggesting that DNA methylation changes may directly contribute to transcriptional alterations during cardiac aging. In contrast, DMRs and DARs that shared the same neighboring gene did not exhibit significant correlation (Supp. Fig. 11C), indicating that DNA methylation and chromatin accessibility may act through largely distinct mechanisms. These findings support a model in which age-associated DNA hypermethylation contributes to gene expression changes.

To directly investigate the functional consequences of age-associated DNA hypermethylation in CMs, we employed a cardiotropic MyoAAV4E vector derived from AAV9^52^ to overexpress the DNA methyltransferase DNMT3A, the predominant DNA methyltransferase isoform in CMs (Supp. Fig. 11D), under the control of the CM-specific *Tnnt2* promoter in young adult mouse hearts (Fig. 4E-F). Age-matched control mice were injected with an MyoAAV4E vector expressing GFP under the same promoter. Successful *Dnmt3a* overexpression (OE) was confirmed by RTqPCR (Supp. Fig. 11E), Western blot (Fig. 4G), and immunohistochemistry (Fig. 4H).

To assess the impact of *Dnmt3a* OE on cardiac function over time, we performed echocardiography every month for 3 months. Systolic function, as measured by ejection fraction (EF), was initially reduced in *Dnmt3a* OE mice, but it recovered to baseline levels by months 2 and 3 (Fig. 4I). In contrast, diastolic function, assessed by the E/A ratio, declined in *Dnmt3a* OE mice throughout the 3-month period (Fig. 4J). *Dnmt3a* OE mice also developed cardiac hypertrophy, as evidenced by increased heart weight to body weight ratio compared to controls at the end of the 3-month period (Fig. 4K), recapitulating another key feature of the aged heart phenotype.

We used RRBS to assess the effect of DNMT3A overexpression on DNA methylation. In CM nuclei from GFP control and *Dnmt3a* OE hearts, global methylation levels at CpG islands remained unchanged following DNMT3A overexpression (Supp. Fig. 11F). However, 2673 regions were hypermethylated in *Dnmt3a* OE compared to control CMs (Supp. Fig. 11G), with a corresponding expansion in the number of genes associated with these hypermethylated regions (Supp. Fig. 11H). Gene ontology analysis of differentially methylated genes revealed enrichment in pathways related to Wnt signaling, DNA-binding repression, ion transport, and phosphatidylinositide-mediated signaling—hallmarks that were also observed in the GO term profiles of methylated genes in aged versus young CMs (Supp. Fig. 11I). 178 genes were hypermethylated in both aged CMs (compared to young) and *Dnmt3a* OE CMs (compared to GFP controls) (Supp. Fig. 11J). This overlap was highly statistically significant (hypergeometric test, p=4.7E-60), indicating that *Dnmt3a* overexpression recapitulates aspects of age-associated DNA hypermethylation. These shared targets were enriched in pathways regulating calcium homeostasis, PI3K/phosphatidylinositide signaling, DNA-binding repression, and cell adhesion (Supp. Fig. 11K), suggesting that these may represent convergent downstream consequences of increased methylation, whether driven by aging or DNMT3A overexpression.

RNA-seq analysis of ventricular myocardium identified 417 DEGs (216 up and 201 down) in *Dnmt3a* OE hearts compared to controls (Fig. 4L). Gene ontology analysis highlighted significant alterations in pathways related to calcium and ion handling (Fig. 4M-N), overlapping with calcium homeostasis pathways identified in our RRBS analysis and suggesting a convergent effect of DNMT3A-mediated hypermethylation on calcium regulation. Other notable overlapping pathways were muscle function and organization of cell junction. Importantly, *Dnmt3a* OE significantly upregulated the CM aging marker *Gm12381* (Fig. 4O), further supporting the role of DNA hypermethylation in driving age-related CM dysfunction.

### Esrrg regulates age-related diastolic dysfunction

In our multi-omics data, we observed widespread changes in RNA transcription as well as pathways related to transcription factor binding in CMs (Fig 3I, Supp. Fig. 8C, and 8H-I). Therefore, we sought to identify transcription factors that regulate CM aging and age-related dysfunction. Using our concurrent single-nucleus ATAC and RNA data, we mapped transcription factors and their target enhancers and genes, collectively referred to as eRegulons, using SCENIC+^53^ (Fig. 5A, Table S5). eRegulon activity successfully separated the major cardiac cell types (Supp. Fig. 12A) and separated old vs young CMs (Supp. Fig. 12B). We identified TFs associated with different cell types and cell states (Supp. Fig. 12C). To prioritize TFs potentially involved in cardiac aging, we developed a dual scoring system. First, we calculated the proportion of age-related genes among all targets for each TF (age-related target ratio). Second, we incorporated the importance scores derived from SCENIC+ for interactions between TFs and their age-related target genes. Both metrics were converted to Z-scores and combined to generate a comprehensive ranking score. Based on this scoring system, ESRRG (estrogen-related receptor gamma) emerged as the top candidate (Supp. Fig. 12D). ESSRG has recently been recognized as a key driver of CM maturation, gene expression, and ventricular CM identity^54–57^. Moreover, six different regions near *Esrrg*, including the promoter, were hypermethylated in aged CMs (Fig. 5B), placing it in the top 1.5% most hyper methylated genes in our RRBS dataset (Supp. Fig. 12E). *Esrrg* was also one of the few downregulated genes in aged CMs (Fig. 5C). These data suggested that *Esrrg* hypermethylation and decreased expression could be directly involved in the emergence of age-related CM dysfunction.

**Figure 5.**
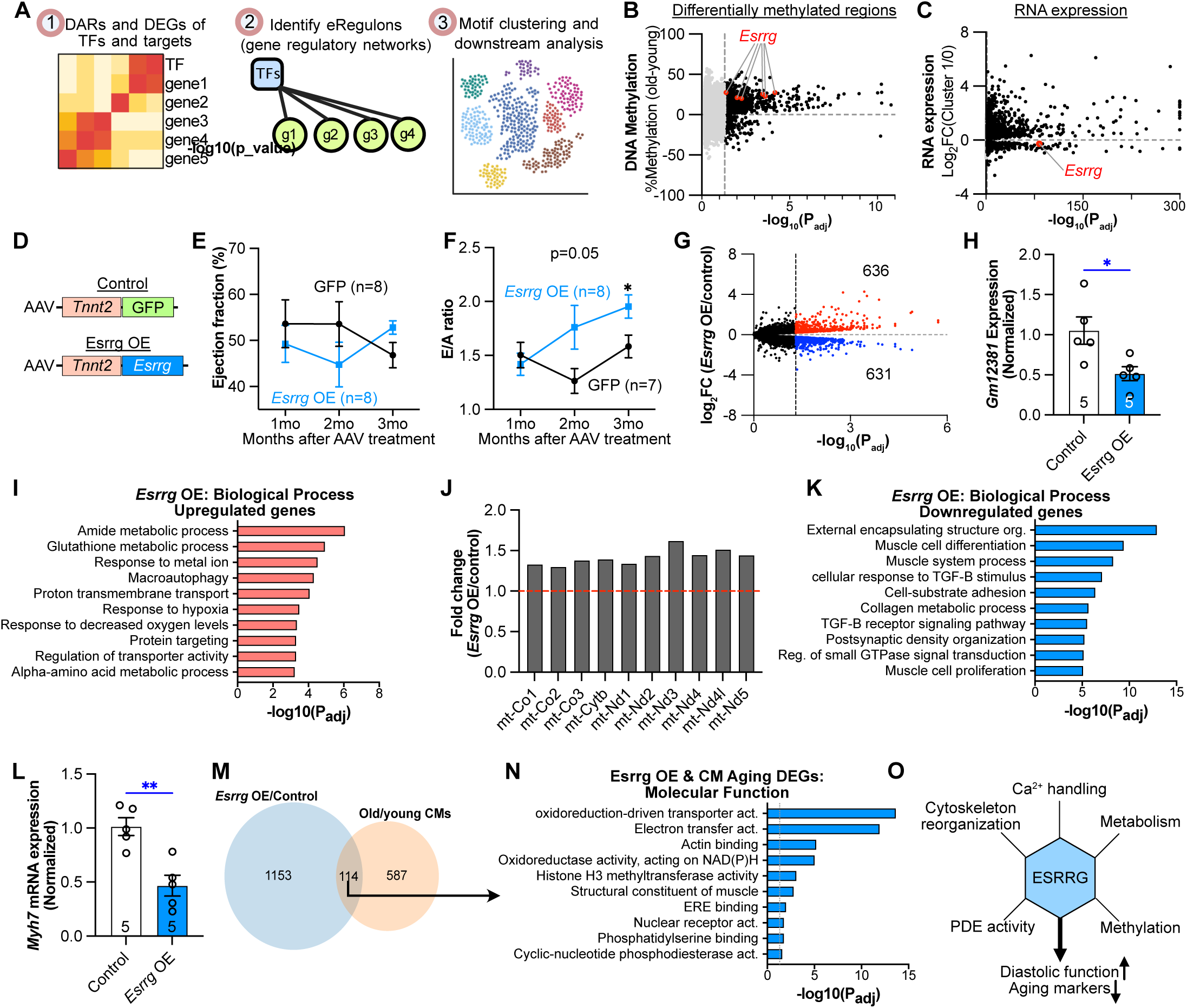
*Esrrg* regulates age-related diastolic dysfunction. **(A)** Schematic of the SCENIC+ pipeline for gene regulatory network analysis. **(B)** Age-related change in CM DNA methylation. Regions near *Esrrg* are highlighted in red. **(C)** Age-related changes in gene expression, with *Esrrg* highlighted in red. **(D)** Schematic representation of AAV constructs for *Esrrg* overexpression (OE). Constructs were injected into adult mice. **(E-F)** Ejection fraction (EF) and E/A ratio of *Esrrg* OE and control mice. *t*-test. **(G)** Volcano plot of RNA-seq data showing differentially expressed genes (DEGs) between *Esrrg* OE and control hearts. **(H)** RTqPCR showing *Gm12381* downregulation in *Esrrg* OE hearts (n=5 per group). *t*-test. **(I)** Biological process gene ontology terms enriched for upregulated DEGs. **(J)** RNA-seq comparison of *Esrrg* OE and control hearts. Mitochondrial genes were upregulated in *Esrrg* OE hearts. **(K)** Biological processes gene ontology terms enriched for downregulated DEGs. **(N)** RTqPCR showing decreased *Myh7* expression in *Esrrg* OE compared to control hearts (n=5 per group). **(M)** Venn diagram showing overlap between DEGs of *Esrrg* OE vs. control and DEGs of old vs. young cardiomyocytes. **(N)** Molecular function GO terms enriched for the common genes identified in M. Summary schematic of *Esrrg* function in cardiac aging. Data are presented as mean ± SEM. *p < 0.05, **p < 0.01.

To investigate the role of *Esrrg* in age-related cardiac dysfunction, we overexpressed *Esrrg* in 3-month-old mouse hearts using a MyoAAV4E vector under the control of the *Tnnt2* promoter (Fig. 5D and Supp. Fig. 12F). While systolic function remained unchanged over the following three months, the E/A ratio continuously increased, indicative of improved diastolic function (Fig. 5E-F). To determine the molecular pathways related to the *Esrrg*-driven improvement in diastolic function, we performed RNA sequencing on *Esrrg*-overexpressing (*Esrrg* OE) and control hearts. We identified over 1,267 genes that were significantly differentially expressed between *Esrrg* OE and control mice (Fig. 5G). Notably, the level of CM aging marker *Gm12381* was lower in *Esrrg* OE compared to control, highlighting its age-reversing transcriptional effect (Fig. 5H). Gene ontology analysis of the upregulated DEGs revealed biological processes such as metabolism and oxidative response that were changed by both aging and *Esrrg* overexpression (Fig. 5I). Notably, autophagy, a key process commonly dysregulated during aging, was among the enriched GO terms associated with upregulated genes in the ESRRG OE model. Consistent with *Esrrg*’s known role in promoting mitochondrial biogenesis, *Esrrg* overexpression increased mitochondrial gene expression (Fig. 5J). In contrast, downregulated genes were mainly related to ECM organization and TGF-beta signaling (Fig. 5K). The cardiac stress marker *Myh7* was also downregulated (Fig. 5L). In terms of molecular function, upregulated genes were related to ribosome biosynthesis and metabolism while downregulated genes were mainly related to ubiquitin ligase activity, ECM remodeling, regulation of histone methylation and cell adhesion (Supp. Fig. 12G-H).

We then compared the DEGs found in *Esrrg* OE with those found in CM aging (Fig. 5M). The 114 genes common between the two lists were associated with pathways such as actinin binding, histone methylation, oxidative response and PDE activity, which were all also on the list of top affected pathways between old and young CMs (Fig. 5N and Supp. Fig. 12G-H).

Next, we tested whether *Esrrg* overexpression is sufficient to reverse aging phenotypes. We treated 24-month-old mice with AAV-Essrrg and measured the effect on cardiac size and function. Four months after injection, *Esrrg OE* did not alter systolic function, diastolic function, cardiac hypertrophy, or expression of aging marker Gm12381 (Supp. Fig. 13A-E), indicating that these age-driven phenotypes are more complex and cannot be reversed by ESRRG overexpression alone.

Finally, we investigated whether ESRRG deletion could produce the inverse phenotype of ESRRG overexpression by inducing diastolic dysfunction. We employed a CRISPR/Cas9-based approach, delivering AAV-Tnnt2-Cre-U6-gRNA AAV vectors containing five different gRNAs targeting *Esrrg* to ensure efficient knockout. In this system, Cre recombinase activates Cas9 expression from Rosa26^fsCas9^ mice by excising a loxP-flanked stop cassette, and the gRNAs direct Cas9 inactivate *Esrrg*. Contrary to our hypothesis, *Esrrg* knockout caused severe systolic dysfunction and death (Supp. Fig. 13F-G). This finding suggests that ESRRG is essential for maintaining cardiac contractile function, and its complete ablation causes a more profound cardiac phenotype than observed with partial reduction observed during normal aging.

Our findings demonstrate that cardiac-specific ESRRG overexpression in adult mice does not result in systolic dysfunction or overt cardiac toxicity, distinguishing our model from prior studies in which constitutive, developmental ESRRG overexpression led to severe cardiac dysfunction and early lethality^58^. This divergence is likely attributable to the critical role ESRRG plays during cardiac development^56^, where its expression must be tightly regulated to ensure proper maturation and differentiation of cardiomyocytes. In contrast, mature adult cardiomyocytes have completed these developmental processes and are therefore less susceptible to the disruptive effects of ESRRG overexpression. Our results support the notion that the functional consequences of modulating ESRRG are highly context-dependent, with adverse effects manifesting only when homeostatic developmental programs are perturbed. Thus, while ESRRG activation in the adult heart appears safe in the context of our model, strategies targeting ESRRG must consider the developmental timing and cellular context to avoid unintended consequences.

In summary, we identified *Esrrg* as a key regulator of age-related cardiac dysfunction. *Esrrg* overexpression altered pathways related to calcium handling, cytoskeleton organization, metabolism, PDE activity, and histone methylation, leading to improved diastolic function and a reduction in aging markers (Fig. 5O).

## Discussion

In this study, we employed a multi-omics approach combining single-nucleus RNA-seq, ATAC-seq, and RRBS to characterize the cellular and molecular landscape of cardiac aging. Our results reveal cell type-specific transcriptional and epigenetic alterations that underlie age-associated phenotypic changes in the heart, including CM hypertrophy, fibrosis, and diastolic dysfunction. Furthermore, we identify DNA hypermethylation as a key driver of age-related CM dysfunction and *Esrrg* as novel regulator of age-related cardiac dysfunction.

Our multi-omic, cell type-specific approach offers several advances over previous studies of cardiac aging. Prior investigations have predominantly relied on bulk RNA-seq analyses^59–64^, which cannot distinguish between CM-specific changes and those occurring in other cardiac cell types. Due to technical challenges of isolating single CMs for single cell transcriptomics, prior single-cell aging studies predominantly captured non-myocytes^65,66^. Some recent studies have begun to employ single-nucleus approaches, but these have been limited by temporal resolution, sample sizes, and use of either snRNA-seq alone, or with separate rather than concurrent snATAC-seq^67,68^. Furthermore, previous investigations into cardiac epigenetic changes during aging^69^ did not detect the CM-specific hypermethylation phenotype we observed, likely due to critical cell type-specific differences that can only be captured through targeted analyses. Other studies conducted in zebrafish^70–72^ provide valuable comparative data but may not directly translate to mammalian cardiac aging given the significant differences in CM behavior between species. Although one recent study employed snATAC-seq^73^ and observed increased chromatin accessibility similar to our findings, it lacked CM-specific analysis. Our concurrent snRNA-seq and snATAC-seq approach, combined with RRBS and robust validation experiments, provides unprecedented resolution of the molecular landscape of CM aging and enables the identification of regulators of age-related cardiac dysfunction.

Our phenotypic characterization of young and aged hearts confirmed previous findings of age-associated cardiac hypertrophy, fibrosis, and diastolic dysfunction^53,74^. We observed significant heterogeneity in CM aging, with a subpopulation of aged CMs exhibiting marked pathological remodeling characterized by *Myh7* upregulation and localization to fibrotic regions, consistent with prior studies^75^. This heterogeneity underscores the importance of single-cell resolution in dissecting the complex mechanisms of cardiac aging and suggests that targeting specific CM subpopulations may be a promising strategy for age-related heart failure.

snRNA-seq analysis has been used to assess cellular composition. Our snRNA-seq data suggested that CM number declines during cardiac aging, while the number of ECs per CM increased. Our direct validation experiments supported the loss of CMs during aging, which likely contributes to aging phenotypes and increases the vulnerability of the aged heart to injury. However, contrary to the snRNA-seq data, our direct validation experiments indicated that the proportion of ECs to CMs was comparable between young and aged hearts. This inconsistency likely stems from differential efficiency of nuclei isolation from different cell types or aging time points. These technical considerations highlight the importance of validating single-nucleus findings with complementary approaches such as histology, particularly when drawing conclusions about quantitative changes in cellular composition during processes like aging.

Identifying reliable markers of cardiac aging remains challenging for both research and future clinical applications. Our snRNA-seq data revealed *Gm12381* as a lncRNA that selectively increases in CMs during aging. Importantly, *Gm12381* was not induced by isoproterenol-mediated stress, highlighting the lncRNA as a valuable biomarker that can distinguish aging from general stress responses in the heart. However, the functional role of *Gm12381* and other age-associated lncRNAs in cardiac aging remains unexplored. Future studies should investigate whether these lncRNAs play causal roles in age-related cardiac dysfunction or merely serve as markers of the aging process, potentially uncovering novel regulatory mechanisms and therapeutic targets.

Integration of snRNA-seq and snATAC-seq data revealed dramatic epigenetic remodeling in aged CMs, with widespread increases in chromatin accessibility correlating with transcriptional changes. These findings are consistent with recent studies demonstrating a global increase in chromatin accessibility in aging in other cell types and highlight the role of epigenetic dysregulation in driving age-related cellular dysfunction^36,76^.

We observed increased DNA methylation in aged CMs. Although expression of *Dnmt3a*, the main DNA methyltransferase in CMs, was not elevated in aged CMs, we hypothesize that sustained *Dnmt3a* expression across the lifespan leads to progressive accumulation of DNA methylation. Hypermethylation has been observed in diverse cell types during aging, but little is known about its causative contribution to aging phenotypes. We directly tested the causal role of DNA hypermethylation in age-related CM dysfunction by overexpressing *Dnmt3a* in young adult mouse hearts. Remarkably, *Dnmt3a* overexpression recapitulated key features of the aged heart phenotype, including diastolic dysfunction and hypertrophy. These results provide the first direct evidence that DNA hypermethylation is sufficient to drive age-related cardiac decline and identify *Dnmt3a* as a potential therapeutic target for age-related heart failure. Long-term experiments are needed to determine if long term inhibition of DNA methylation, e.g. by *Dnmt3a* inhibition or loss-of-function, is sufficient to mitigate cardiac aging phenotypes. It will also be interesting to examine the contribution of *Dnmt3a* to heart failure with preserved ejection fraction (HFpEF), the predominant form of age-related heart failure^77^.

Unbiased analysis of our single cell multiomic data nominated ESRRG as a key transcription factor mediator of age-related diastolic dysfunction. ESRRG is a nuclear receptor that regulates metabolism^78^, CM maturation^54–56,79^, and ventricular CM identity^57^. Although its DNA-binding and ligand-binding domains are homologous to the estrogen receptor^80^, ESRRG does not bind estrogen and its ligand is currently unknown. ESRRG was among the most downregulated and hypermethylated genes in aged CMs, and its overexpression in young mouse hearts improved diastolic function and downregulated the CM aging marker *Gm12381*. These findings suggest that *Esrrg* may maintain youthful CM function by coordinating multiple aspects of cellular homeostasis, including metabolism, calcium handling, and histone modification. However, ESRRG overexpression alone was not sufficient to reverse cardiac aging phenotypes in aged mice, pointing to the complexity and multifactorial nature of the aging process.

While our study provides novel insights into the molecular basis of cardiac aging, it also raises several questions that warrant further investigation. We identified striking changes in chromatin accessibility, DNA methylation, and lncRNA expression in aged CMs. The mechanisms underlying these changes and their functional consequences require further investigation. The translational relevance of our findings in mice to human cardiac aging remains to be established. Comparative analyses of aging hearts from mice and humans will be important steps in advancing our understanding of human cardiac aging. These studies will be crucial in determining whether the epigenetic and transcriptional changes we observed in mice are conserved in humans and whether targeting *Dnmt3a* could be a viable therapeutic strategy for age-related cardiac dysfunction in humans. Lastly, while our scMultiome dataset offers significant new insights into the mechanisms of cardiac aging, we acknowledge that it includes only male samples. Followup bulk ATAC-seq experiments did not detect sex differences at FDR<0.05, but additional focused studies are required for fully explore the interaction between sex and aging.

In conclusion, this study provides an extensive molecular atlas of murine cardiac aging and identifies DNA hypermethylation and Esrrg downregulation as significant regulators of age-related cardiac phenotypes in mice. These findings not only advance our fundamental understanding of the aging heart but also inform the development of targeted therapies for age-related heart failure.

## Methods

### Mice and tissues

All experiments with mice were performed under protocols approved by the Boston Children’s Hospital Institutional Animal Care and Use Committee. Aged (25-month-old) and young (4-month-old) mice (C57BL/6J) were obtained from Jackson Laboratories. For multi-omic, echocardiography, and pressure-volume (PV) loop experiments, only male mice were used to reduce variability. The remaining experiments were conducted using mixed cohorts of both male and female mice. All mice were maintained on a regular diet. In AAV studies, mice within the same cage were injected with different AAVs to minimize variability between experimental groups. Old and young mouse (C57BL/6JN) tissues and sections were obtained through National Institute of Aging to support this project.

### Histology

7 μm sections were prepared from paraffin-embedded (pre-fixed for 24 h) and OCT-embedded frozen tissues. OCT sections were thawed, washed with PBS, and fixed with 4% paraformaldehyde (PFA) for 15 min. Paraffin sections were treated with xylene and rehydrated through an ethanol gradient.

For immunostaining, sections were blocked (2% BSA, 0.3% Triton X-100 in PBS) for 1 h, incubated with primary antibodies (1:200) overnight at 4°C, washed, and incubated with secondary antibodies/WGA (1:200) and DAPI (1:1000) for 1 h. Sections were mounted using mounting media (Thermo Fisher, P36961). Antibodies and dyes are listed in Table S5.

Sirius Red/Fast Green staining was performed by incubating sections in 0.1% Fast Green (in 1.2% picric acid) for 10 min, washing in 1% acetic acid and tap water, and staining with 0.1% Sirius Red (in picric acid) for 30 min. Sections were dehydrated, cleared in xylene, and mounted in DPX mountant.

### Microscopy

Confocal images were acquired using an Olympus FV3000RS microscope with 30X and 60X oil immersion objectives. Epifluorescence images for quantification of CMs and fibrosis were obtained using a Keyence All-in-One Fluorescence Microscope with a 10X objective.

### RT-qPCR

Total RNA was isolated using the Quick RNA Miniprep Plus Kit (Zymo Research, R1058) and quantified using NanoDrop. High-quality RNA samples (>1.8 230/260 and 260/280 ratios) were converted to cDNA using the iScript cDNA Synthesis Kit (Bio-Rad Laboratories, 1708891) with 1 μg RNA per reaction. cDNA was quantified using SYBR Green (Life Technologies, 4368708) on Bio-Rad CFX96 or CFX384 instruments. Significant outliers based on Grubb’s test were excluded. Gene expression was normalized to the housekeeping gene *Ppia*. Primer sequences are listed in Table S6.

### Western Blotting

Tissues were lysed in RIPA Lysis and Extraction Buffer (Life Technologies, 89900) with 5 mm stainless steel beads (Qiagen, 69989) using a TissueLyser II at 40 Hz for 1 min. Lysates were incubated on ice for 2 h, centrifuged at 13,000 x *g* for 20 min at 4°C, and supernatants were collected. Protein concentration was determined using the Pierce BSA kit (Thermo Fisher, 23225). 30 μg of protein per lane was separated and transferred using the Trans-Blot SD semi-dry transfer system (Bio-Rad Laboratories). 0.45 µm nitrocellulose membranes (Bio-Rad Laboratories, 1620115) were blocked in 5% BSA in 0.1% TBS-T for 1 h at room temperature, incubated with primary antibodies (1:1000) overnight at 4°C, washed twice with 0.1% TBS-T, and incubated with HRP-conjugated secondary antibodies for 1 h at room temperature. After washing, blots were incubated with HRP substrate (Thermo Fisher Scientific, WBKLS0500) for 1 min and imaged. Band intensities were quantified using ImageJ.

### Echocardiography

Mice were anesthetized using 2.5% isoflurane, and echocardiography was performed using the VEVO 3100 system. Fractional shortening was measured using M-mode in the short-axis view to assess systolic function. Diastolic function was evaluated by measuring E/A ratios using pulsed-wave Doppler. Echocardiography was performed blinded to group assignments.

### Image processing and analysis

For WGA images, CellPose^81^ was used to measure the cross-sectional area of CMs. For Sirius Red/Fast Green staining, the color deconvolution function was applied. Built-in ImageJ functions were used for the remaining analyses.

### Invasive hemodynamics

Invasive hemodynamics were performed as previously described^82^. A high-fidelity pressure catheter (1.0 Fr. micro-tip, Transonic Systems Inc) was inserted through the right carotid artery to obtain volume-pressure loops. Data were recorded using a PowerLab system (ADInstruments).

### CM isolation and staining

CMs were isolated following a previously established protocol^37^. Briefly, CMs were fixed in 4% PFA and incubated in a 2 mg/ml collagenase II solution in PBS until the tissue was completely lysed. Isolated CMs were permeabilized using 0.5% Triton X-100 solution and blocked at room temperature for 1 hour in 2% BSA solution. Primary antibodies were applied overnight at 4°C, followed by secondary antibodies for 1 hour at room temperature in a 2% BSA, 0.1% Triton X-100 solution. Stained CMs were either mounted on slides or seeded on black flat-bottom 96-well plates (Sigma Aldrich, M0562-32EA).

### AAV construction, packaging, and isolation

AAV plasmid constructs were generated to express *Dnmt3a* (RefSeq: NM_175629.2), *Esrrg* (RefSeq: NM_001243509.1), or GFP under the control of the cardiac-specific *Tnnt2* promoter. The cDNAs were PCR-amplified and cloned into the AAV expression vector downstream of the *Tnnt2* promoter using In-Fusion cloning.

AAV packaging was performed as previously described^83^. Briefly, a mixture of expression and Rep/Cap helper vectors were transfected into HEK 293T cells. Viruses were then purified using an iodixanol density gradient with OptiPrep. For all mice, the MyoAAV-4E^52^ capsid was used. A dose of 2×10^10^ vg/g per mouse was administered through retro-orbital injection into adult mice.

### Bulk RNA-sequencing

Total RNA was isolated using the Quick RNA Miniprep Plus Kit (Zymo Research, R1058), and RNA quality was assessed using the Agilent TapeStation. All samples had RIN values above 8. Sequencing was performed on an Illumina NovaSeq (PE150, 60 M reads per sample). The obtained FASTQ files were processed using the BioJupies platform^84^, and GO terms were identified using the clusterProfiler package ^85^ and further processed with REVIGO ^86^ to remove redundancy. Terms with a dispensability score greater than 0.3 were excluded from the analysis.

### Bulk ATAC sequencing

Bulk-ATAC sequencing was performed using ATAC-Seq kit (Active Motif North America, 53150, based on manufacturer’s instructions. The raw files were processed in encode ATAC-seq pipeline^87^. Reads were trimmed, aligned using Bowtie2, and filtered. Differentially accessibility analyses were done using different normalization methods as previously described^87,88^.

### scMulti-omics

Frozen tissues were transferred from −80°C storage and immediately placed in homogenization buffer (HB; 250 mM Sucrose, 25 mM KCl, 5 mM MgCl_2_, 1 μM DTT, 0.1% Triton-X-100, 10 mM Tris, pH 8.0 supplemented with RNAse inhibitors) solution. Tissues were cut into 3-5 mm pieces on ice, placed in HB solution, and lysed using a TissueLyser II by homogenizing with 5 mm stainless steel beads. Tissues were further incubated on ice for 5 minutes. Nuclei were filtered through a 40 μm cell strainer and centrifuged at 500 g for 5 minutes at 4°C. Pellets were resuspended in storage buffer (SB; 4% BSA in RNAse free PBS supplemented with RNAse inhibitors). 7-AAD (Life Technologies, A1310) was added to the nuclei solutions at a 1:1000 ratio. A small aliquot was taken as a negative control. Nuclei were sort-purified using a BD FACSAria II, and approximately 200,000 nuclei were captured.

Following sorting, nuclei were counted using a Countess 3. 10,000 nuclei per sample were processed using the 10x Genomics Next GEM Single Cell Multiome ATAC + Gene Expression Reagents Kit. Resulting libraries were analyzed using TapeStation and sequenced on an Illumina NovaSeq (PE150, 200M PE reads per sample). Tissues from both timepoints (4- and 28-month-old) were processed simultaneously to prevent batch effects.

### Multi-omics analysis

Fastq files from the 10X Genomics Multiome ATAC + Gene Expression assay were processed using the CellRanger-ARC software (10X Genomics) to generate gene expression and ATAC output files. Further analysis was performed using the Seurat^89^ and Signac^90^ packages in R^90,91^. Quality control was performed by filtering out cells with less than 800 RNA counts, less than 500 RNA features, or more than 10% mitochondrial DNA content. The data was then normalized, and variable features were identified. For the ATAC data, a Chromatin Assay was created using the ATAC counts. The counts were annotated based on the mm10 genome assembly using the Ensembl database (EnsDb.Mmusculus.v79). To identify and remove potential doublets, the DoubletFinder^92^ algorithm was applied. A parameter sweep was performed to determine the optimal pK value (0.005). The homotypic doublet proportion was estimated, and the expected number of doublets was adjusted accordingly. Cells identified as doublets were removed from the dataset.

The RNA and ATAC data were then integrated using the weighted nearest neighbor (WNN) analysis. The RNA data was processed using the SCTransform function, followed by PCA and UMAP (dims = 1:50). For the ATAC data, TF-IDF normalization was applied, followed by singular value decomposition (SVD) and UMAP (dims = 2:50). The multi-modal neighbors were identified using both PCA (dims = 1:50) and LSI (dims = 2:50) reductions.

Differential gene expression and chromatin accessibility analyses were performed using the FindMarkers function in Seurat, with the negative binomial test to identify significant differences between clusters. To associate genes with accessible chromatin regions, the ClosestFeature function from the Signac package was used. GO-Term analyses were done using EnrichR^93^.

Permutation analysis was using the regioneR^94^ package in R. CpG island annotations were obtained from the UCSC Genome Browser database (mm10 assembly). The analysis was restricted to canonical chromosomes, and DAR positions were randomly shuffled across the genome for 1,000 iterations while maintaining their size distribution and chromosome assignments. For each permutation, overlaps between randomized DARs and CpG islands were calculated. The observed number of overlaps was compared to the permuted distribution to assess statistical significance, with fold enrichment calculated as the ratio of observed to mean permuted overlaps.

### Scenic+ analysis

Scenic+^53^ analysis was performed following the documentation with minor modifications. The analysis pipeline was used to infer gene regulatory networks and identify cell state-specific regulons from single-cell RNA-seq data. Transcription factor to gene analyses were done using gradient boosting.To identify transcription factors most relevant to CM aging, we integrated differential expression analysis results (Table S1) with the Scenic+ output (Table S5).

Specifically, we identified the top 1000 differentially expressed genes (excluding mitochondrial genes) and analyzed how they intersected with the predicted transcription factor target genes. We developed a scoring system based on two metrics: (1) the percentage of a transcription factor’s target genes that overlapped with the differentially expressed gene set, and (2) mean importance scores of these overlapping target genes derived from the gradient boosting model. Both metrics were converted to z-scores to account for varying regulon sizes and importance score distributions. The final transcription factor ranking was determined by averaging these z-scores, allowing us to identify transcription factors that both extensively and strongly regulated the differentially expressed genes.

### RRBS sequencing

For RRBS sequencing, the Zymo-Seq RRBS Library Kit (VWR, 76407-310) was used according to the manufacturer’s instructions. 250 ng of DNA was used per sample, and sequencing was performed on an Illumina NovaSeq (PE150, 40 M reads per sample). The obtained FASTQ files were mapped using the mm10 reference genome and analyzed using Bismark^95^ to extract methylation information.

### mtDNA mutation analysis

mtDNA analysis was performed using the mitochondrial genome analysis toolkit (mgatk) on snRNA-seq reads^96^. For each sample, the input BAM file was sorted, and duplicates were marked. Variant calling was conducted using mgatk’s call function with the mm10 mitochondrial reference genome and base quality threshold of 20. Pileup files were generated using Samtools mpileup with appropriate quality filters and a maximum depth setting of 1,000,000. Variant allele frequencies were calculated for each position, and mutation rates were compared between young and aged samples using standard statistical methods.

### Statistics and reproducibility

Unless otherwise noted, results are presented as mean ± standard error of the mean (SEM). Each data point in the graphs represents a unique biological replicate. Statistical analyses were performed in GraphPad Prism or R. For RTqPCR data in Supp. Fig. 3, outliers defined by Grubb’s test were removed.

## Availability of Materials

Materials not available through public repositories or commercial vendors will be provided by the corresponding author upon reasonable request.

## Data access

High-throughput data used in the manuscript are avaialble from the Gene Expression Omnibus, accession numbers <u>GSE281772</u> for scMulti-omics, <u>GSE281602</u> and GSE299673 for RRBS, <u>GSE281775</u> for bulk-RNA sequencing. The reviewer token is kpkzuukgvnyhjcr and opivucyqhvovhad for RRBS, wrgxioaazhotbgb for sc multi-omics, and ehghkyayphwjfgr for bulk RNA-seq datasets.

## Acknowledgments

This research was made possible in part using biomaterials from the NIA Aged Rodent Tissue Bank (Aged Rodent Tissue Bank | National Institute on Aging (nih.gov)) at the University of Washington, Seattle under contractual agreement with the National Institute on Aging (NIA).

## Author contributions

D.Y. conceived the project, designed and performed experiments, analyzed data, and wrote the manuscript. M.A.T. assisted with scMulti-ome experiment. Z.W. contributed to echocardiography analysis. Q.K. and P.K. performed intracardiac pressure measurements. W.T.P. supervised the project, provided guidance on experimental design and data interpretation, and edited the manuscript. All authors reviewed and approved the final version of the manuscript.

