## Supplementary Figure for "Molecular Mechanisms of Cardiomyocyte Aging"

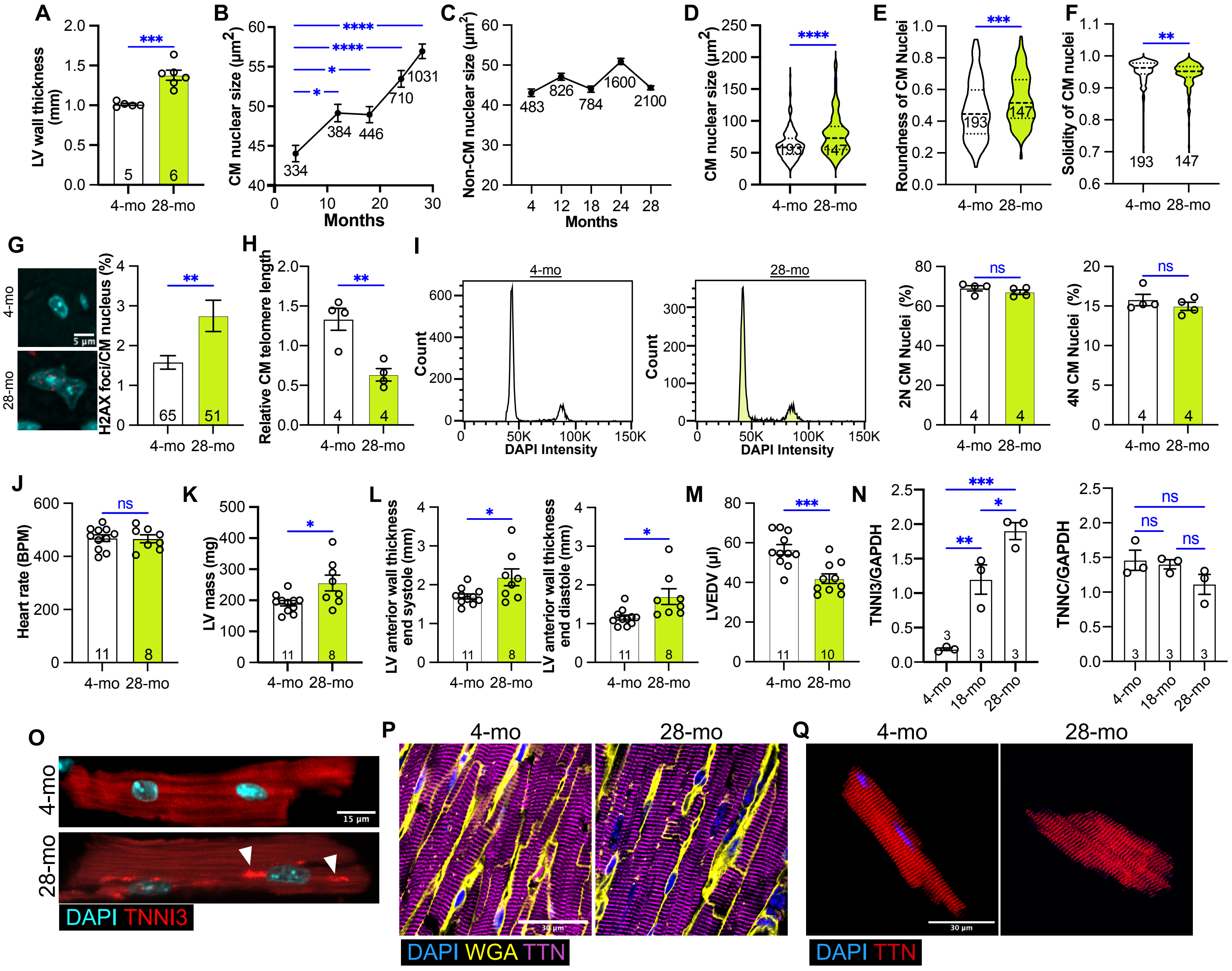


**Supplementary Figure 1. Additional age-associated changes in cardiac structure and function. (A)** Left ventricular wall thickness measured from 4- and 28-month-old heart sections. t-test. **(B-C)** CM (B) and non-CM (C) nuclear size in heart sections at 4, 12, 18, 24, and 28 months. One way ANOVA with Dunnett's post hoc test. n indicates number of nuclei measured. **(D)** CM nuclear size quantification of isolated CMs. *t*-test. **(E)** Roundness of nuclei measured in isolated CMs . t-test. **(F)** Solidity of nuclei measured in isolated CMs. Solidity, defined by area of nuclei by its convex hull area, is a measure of the regularity of the nuclear boundary. *t*-test. **(G)** Immunofluorescence staining for yH2AX foci, a marker of DNA double-strand breaks. Left: Representative images of yH2AX staining in CM nuclei from young and aged hearts. Right: Quantification of yH2AX foci per nucleus in young and aged CMs. *t*-test. **(H)** Relative CM telomere length across age groups. *t*-test. **(I)** CM nuclei ploidy analysis. Left: Representative flow cytometry histograms for 4- and 28-month-old samples. Right: Quantification of 2N and 4N populations. t-test. **(J-M)** Echocardiographic measurement of heart rate (J), left ventricular (LV) mass (K); anterior wall thickness at the end of systole (L, left) and diastole (L, right); and left ventricular end diastolic volume (LVEDV) (M) in young and aged hearts. *t*-test. **(N)** Western blot quantification. ANOVA. (**O**) Representative immunofluorescence images showing TNNI3 localization in isolated CMs from 4-month-old and 28-month-old hearts. Scale bar, 15 μm. (**P-Q**) Immunostaining for titin showing normal sarcomeric organization and localization in cardiac tissue sections (P) and isolated CMs (Q) from young and aged hearts. Scale bar, 30 μm. Data are presented as mean ± SEM for bar graphs and median (solid lines) with quartiles (dashed lines) for violin plots. Numbers in graphs indicate sample size. *p < 0.05, **p < 0.01, ***p < 0.001, ns=not significant.

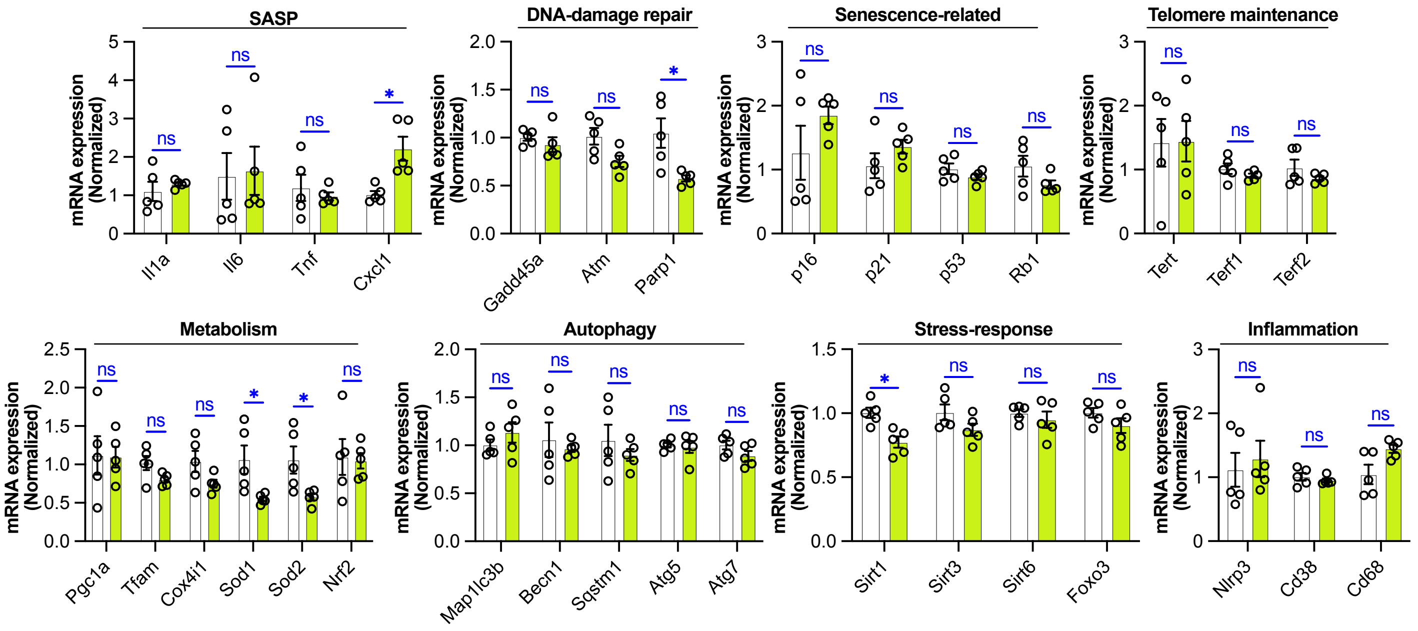


**Supplementary Figure 2. Expression of general aging markers in cardiac aging.** RTqPCR analysis of transcripts associated with senescence-associated secretory phenotype (SASP), DNA damage repair, cellular senescence, telomere maintenance, metabolism, autophagy, stress response, and inflammation in cardiac ventricles from young (4-month-old) and aged (28-month-old) mice. Only *Sod1*, *Sod2*, *Cxcl1*, *Parp1*, and *Sirt1* showed statistically significant differences between age groups. Data are presented as mean ± SEM. n=5 independent biological replicates per group. *$p < 0.05$, ns=not significant.

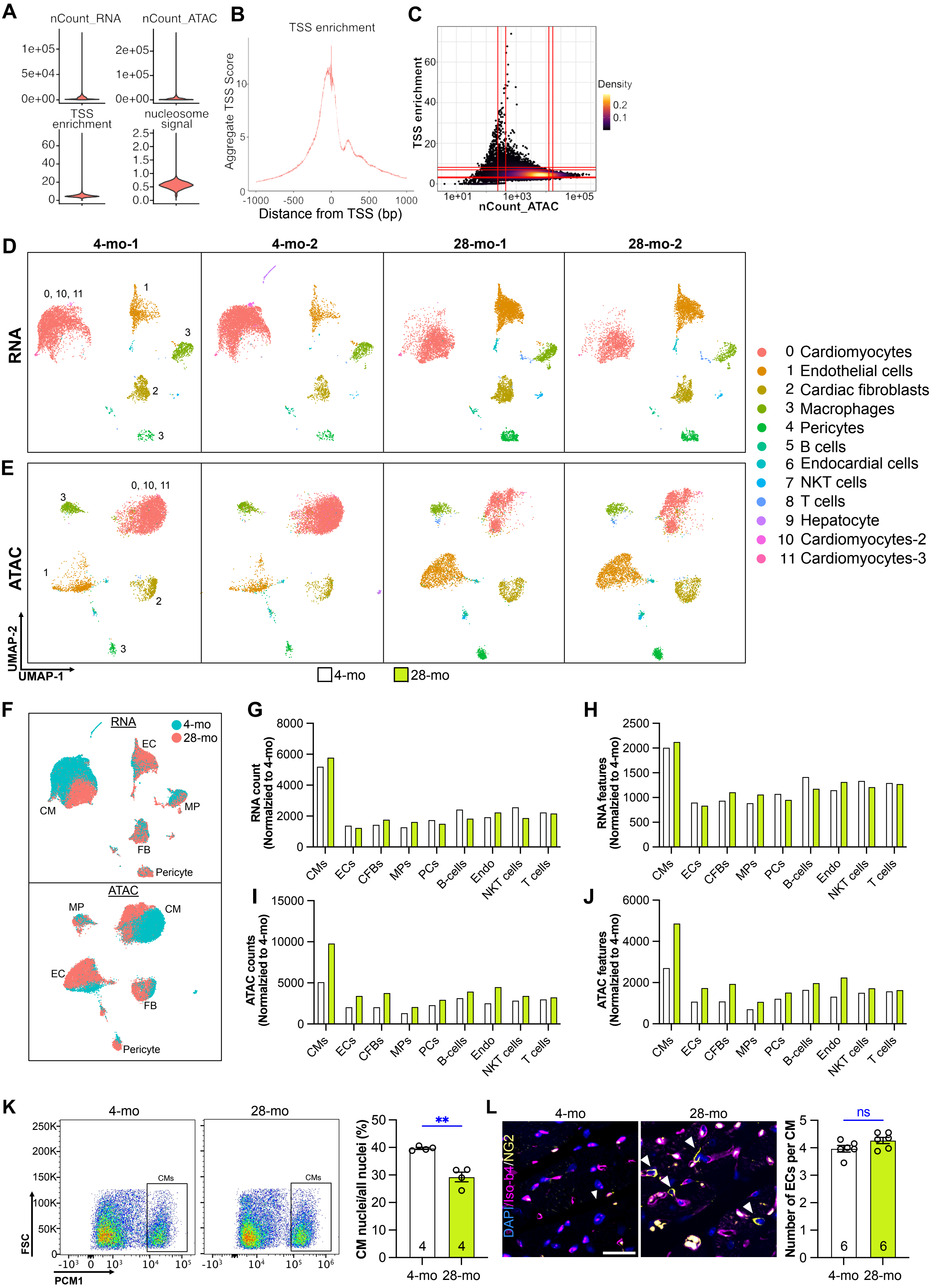


**Supplementary Figure 2**

**Supplementary Figure 3. Concurrent single-nucleus RNA-seq and ATAC-seq data on young and old hearts. (A)** Violin plots showing the distribution of RNA counts, ATAC counts, TSS enrichment scores, and nucleosome signals across samples. **(B)** Average TSS enrichment plot of snATACseq data demonstrating a mean enrichment score of 10. **(C)** Scatter plot of TSS enrichment scores versus ATAC counts for snATACseq data, showing the relationship between sequencing depth and signal quality. Red lines represent 5, 10, 90 and 95% quantiles. **(D-E)** UMAP plots showing individual biological replicates for (A) RNA-seq and (B) ATAC-seq corresponding to the integrated analysis presented in Figure 2B. Each column represents a separate biological replicate. Nuclei and their clustering were consistent across biological replicates. **(F)** UMAP plots of RNA-seq and ATAC-seq data labeled by sample ages, demonstrating clear separation between young and aged samples within specific clusters (e.g. CMs and ECs). **(G)** Per nucleus RNA-seq read count distribution across different cell clusters and age groups. **(H)** Per nucleus number of RNA features (genes) detected across different cell clusters and age groups. **(I)** Per nucleus ATAC-seq read count distribution across different cell clusters and age groups. **(J)** Per nucleus ATAC features (accessible regions) detected across different cell clusters and age groups. **(K)** Flow cytometry analysis of $PCM1^+$ nuclei. Left: Representative flow cytometry plots showing forward scatter (FSC) versus PCM1 staining for isolation of CM nuclei. Right: Quantification of the percentage of $PCM1^+$ (CM) nuclei relative to total cardiac nuclei in young and aged hearts. *t*-test. **(L)** Quantification of cardiac vasculature. Left: Representative images of cardiac sections from 4- and 28-month-old hearts stained with Isolectin B4 (endothelial cells) and NG2 (pericytes). Scale bar, 25 µm. Right: Quantification of Isolectin $B4^+$ and $NG2^+$ cells per CM in young vs old hearts. *t*-test. Data are presented as mean ± SEM $^{**}p < 0.01$, ns=not significant.

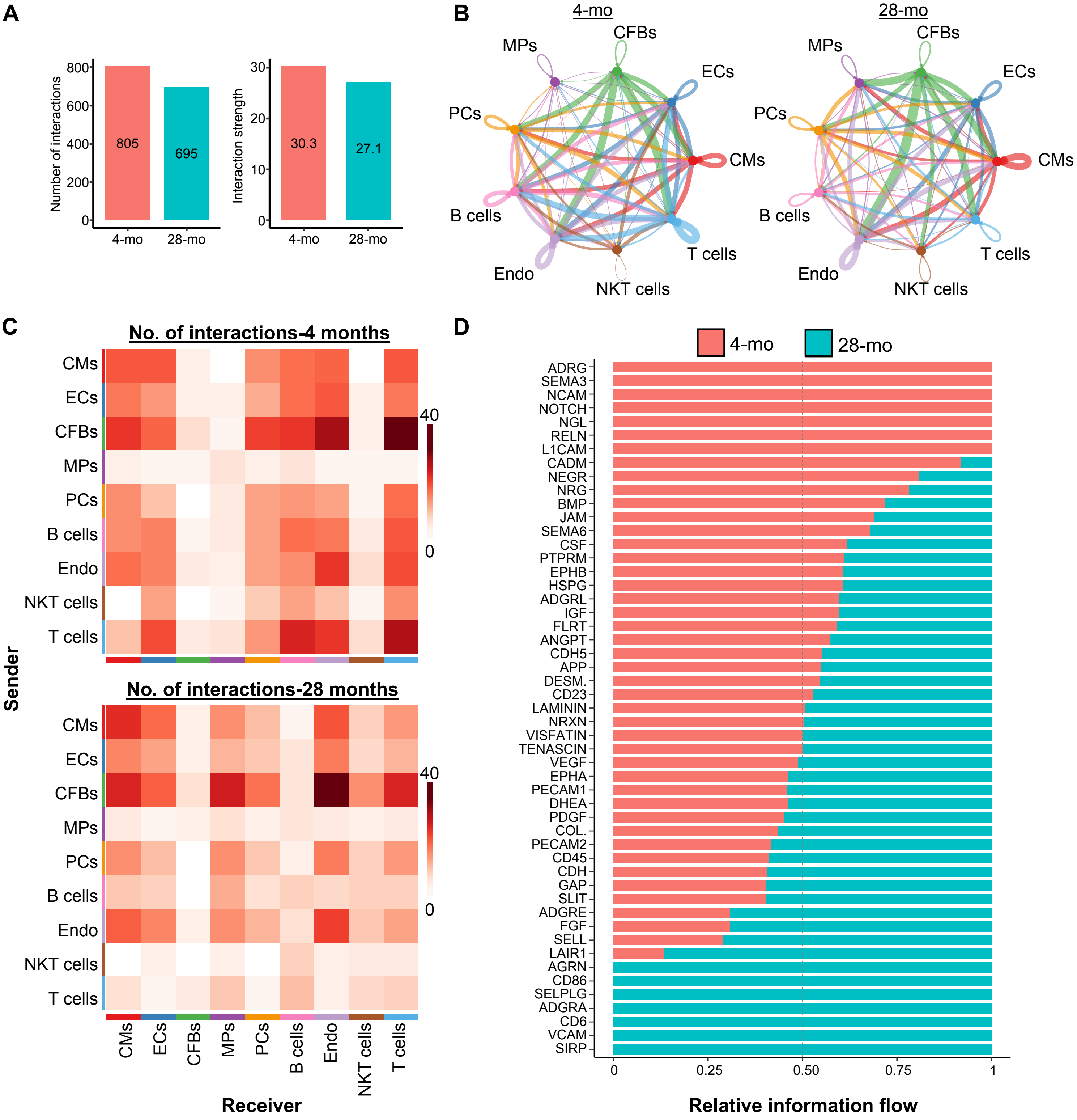


**Supplementary Figure 4. Age-related changes in intercellular communication networks in the heart. (A)** Global comparison of intercellular communications between young and aged hearts, showing reduced interaction frequency and strength in aged cardiac tissue. **(B)** Cell-cell communication patterns in young and aged hearts. **(C)** Heatmap of intercellular communications across all cardiac cell types. Color scale represents the number of interactions. **(D)** Pathways enriched in intercellular communications between young and aged cardiac cell types. Leukocyte proliferation pathways were enriched in aged hearts.

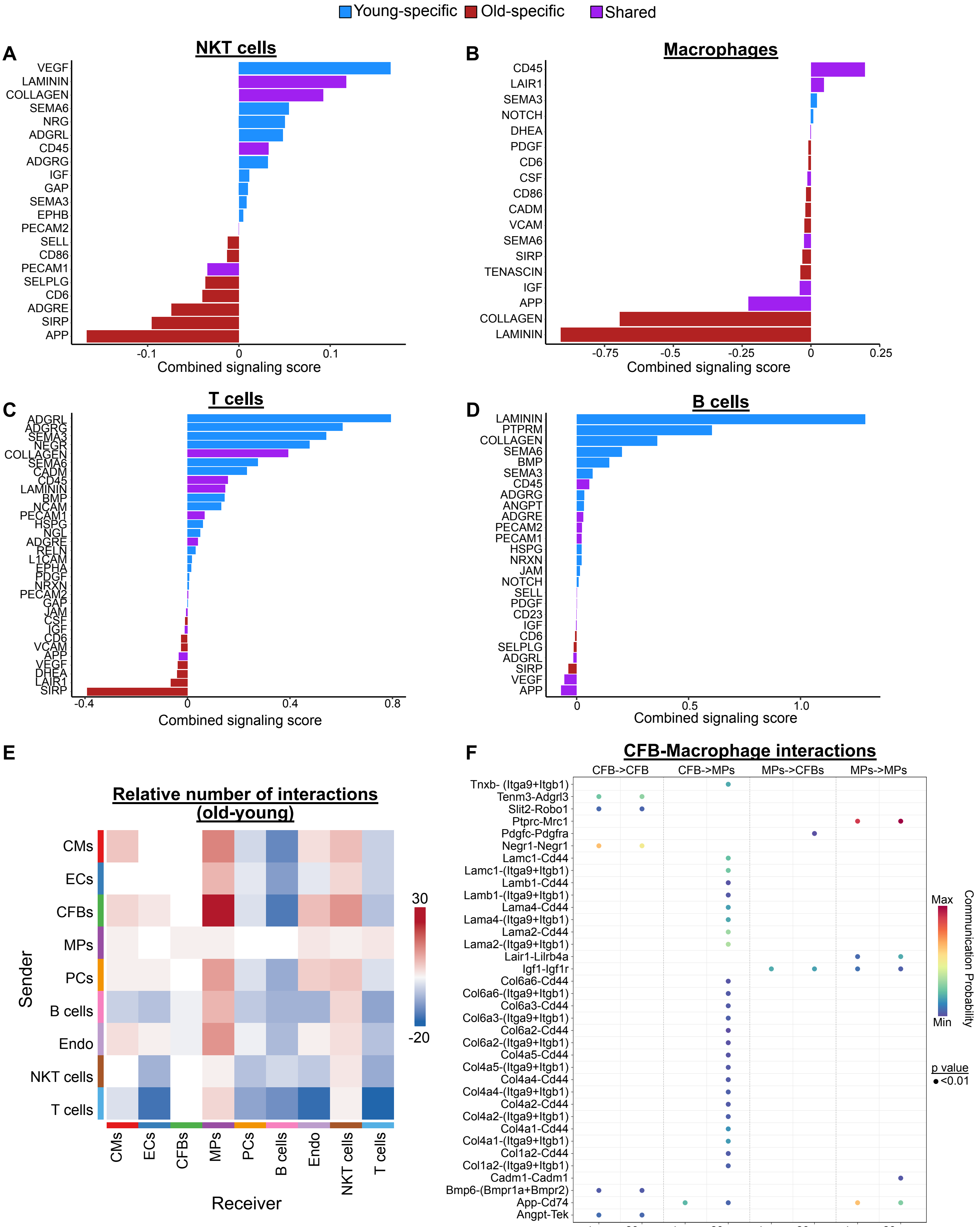


**Supplementary Figure 5.**

**Supplementary Figure 5. Cell type-specific alterations in intercellular signaling during cardiac aging. (A-D)** Analysis of communication networks for indicated cell types, showing differential ligand-receptor utilization between young and aged hearts. **(E)** Cell-type pairs with the most pronounced changes in interaction strength during aging. **(F)** Detailed analysis of fibroblast-macrophage signaling, highlighting enhanced collagen and laminin signaling via ITGA9 and ITGB1 receptors in aged hearts.

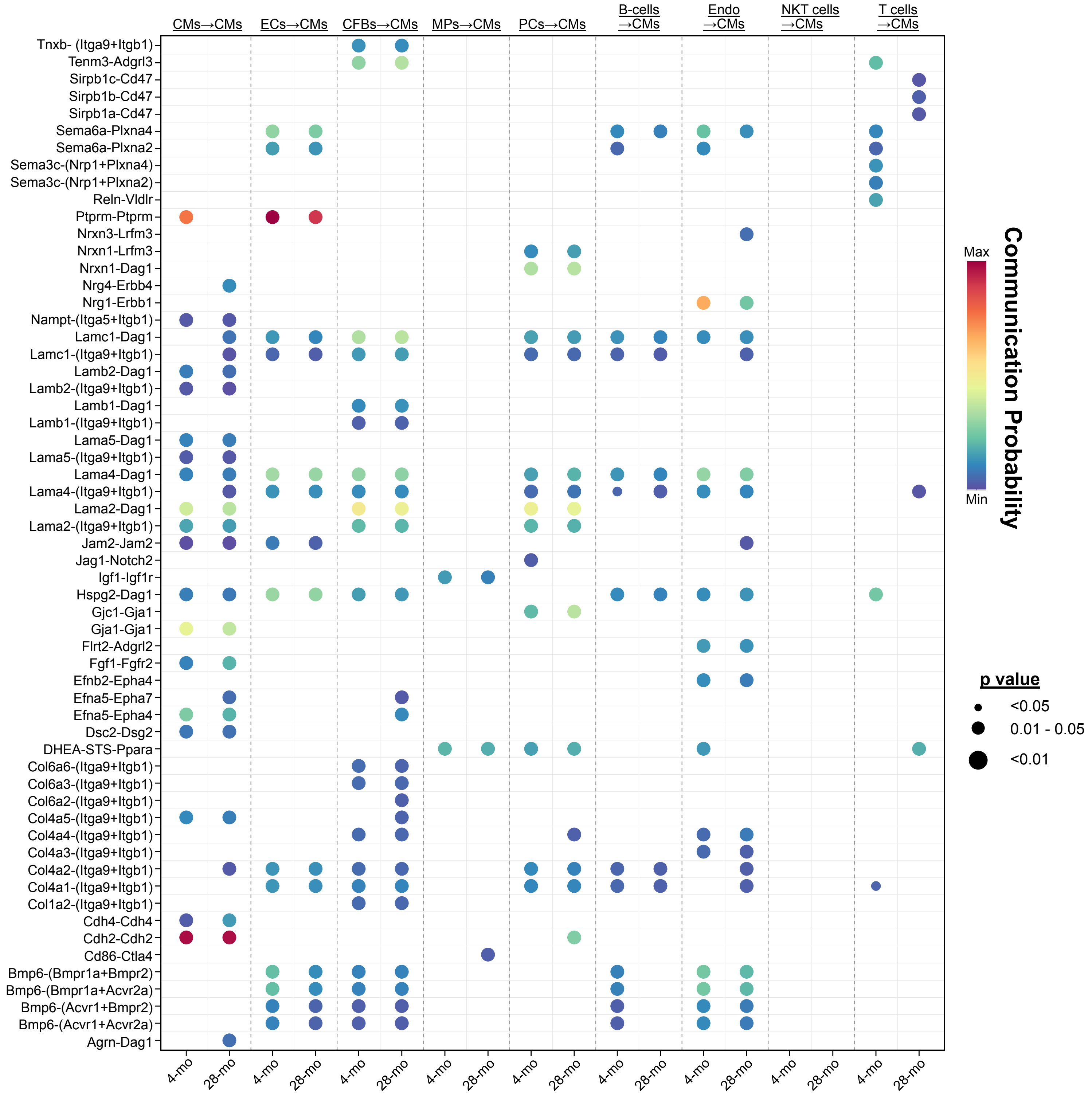


**Supplementary Figure 6. Age-associated alterations in CM intercellular communication.** Analysis of intercellular signaling pathways involving CMs in young versus aged hearts. The diagram highlights changes in direct cell-cell communication from CMs, B cells, and T cells to CMs with aging, while the majority of incoming signaling pathways to CMs remain relatively unchanged.

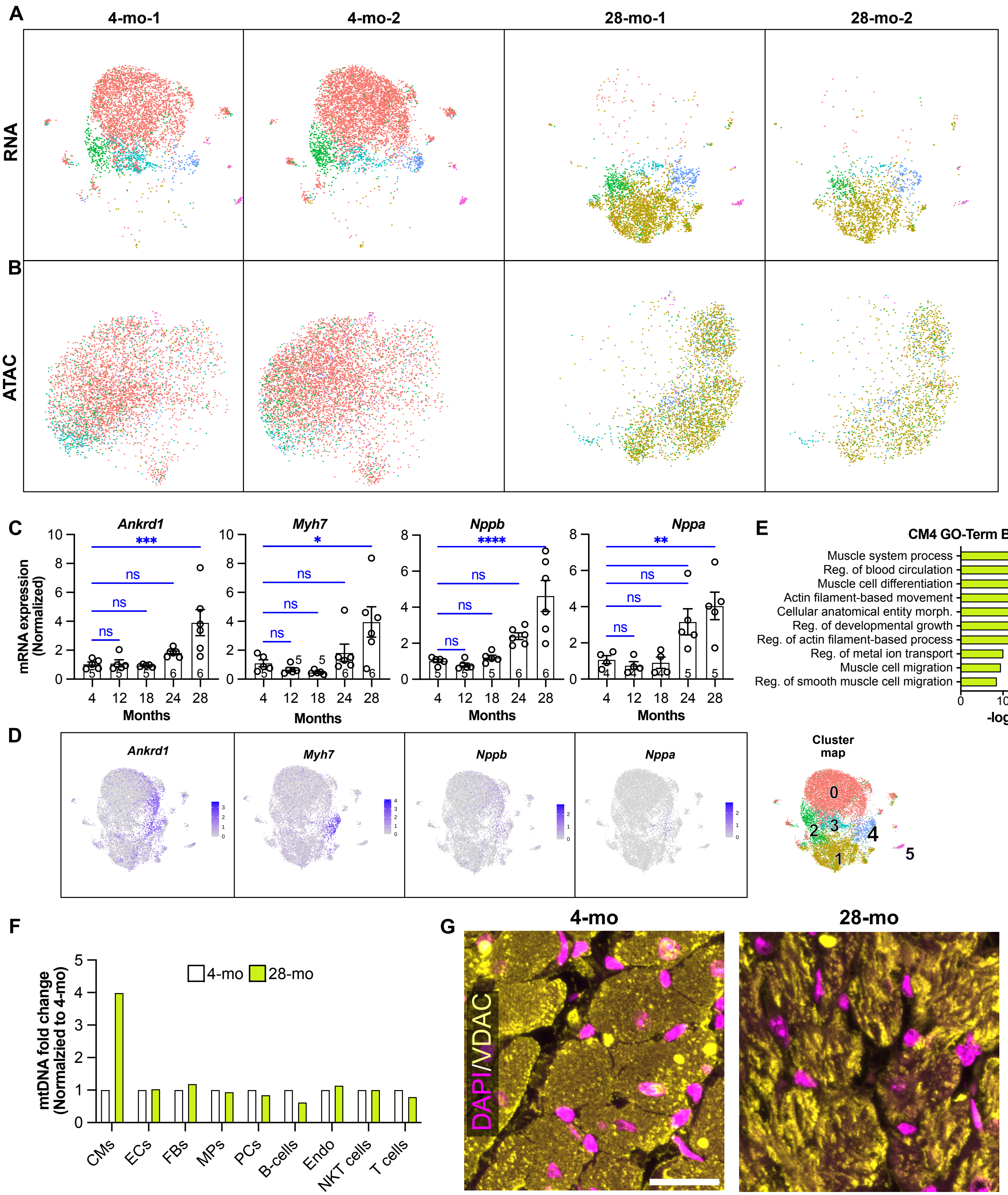


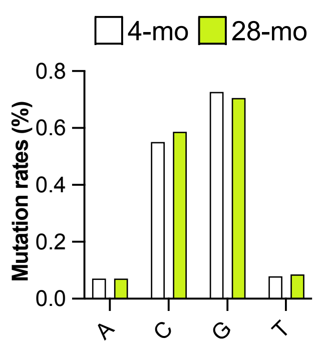


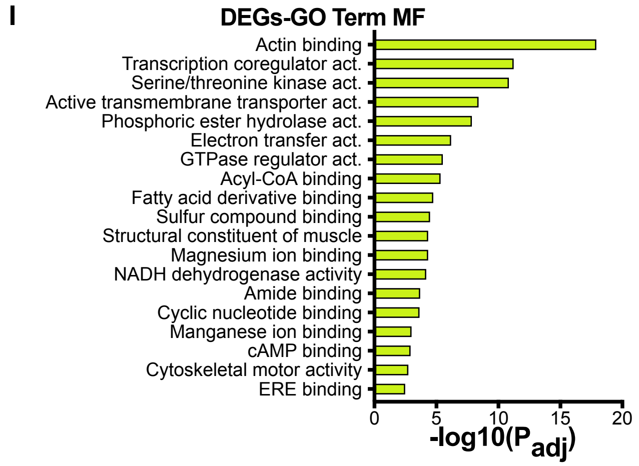


**Supplementary Figure 7.**

**Supplementary Figure 7. Characterization of CM subclusters. (A-B)** snRNAseq and snATACseq UMAP plots showing individual biological replicates for RNA-seq data of subclustered CMs. Each plot represents a separate biological replicate, demonstrating the consistency of CM subcluster identification across samples. **(C)** RTqPCR analysis of CM stress markers (*Ankrd1*, *Myh7, Nppb*, and *Nppa*) in whole heart samples across different age groups (4, 12, 18, 24, and 28 months). One-way ANOVA with Tukey's post-hoc test. **(D)** Feature plots showing expression of stress markers *Ankrd1*, *Myh7, Nppb*, and *Nppa,* highlighting their increased expression in cluster 4. **(E)** Biological process GO terms enriched in cluster 4. **(F)** Mitochondrial DNA (mtDNA) content across different cell clusters and age groups, mtDNA was higher in aged compared to young CMs. **(G)** Representative images of VDAC-1 (voltage-dependent anion channel-1) immunostaining in young and old cardiomyocytes, demonstrating mitochondrial clustering in aged cells. Scale bar, 25 µm. **(H)** Mitochondrial DNA mutation rates comparing 4-month-old and 28-month-old CMs. **(I)** Gene ontology analysis of molecular functions (MF) for differentially expressed genes (DEGs) between young and aged CMs. Data are presented as mean ± SEM. *$p < 0.05$, **$p < 0.01$, ***$p < 0.001$, ****$p<0.0001$, ns=not significant.

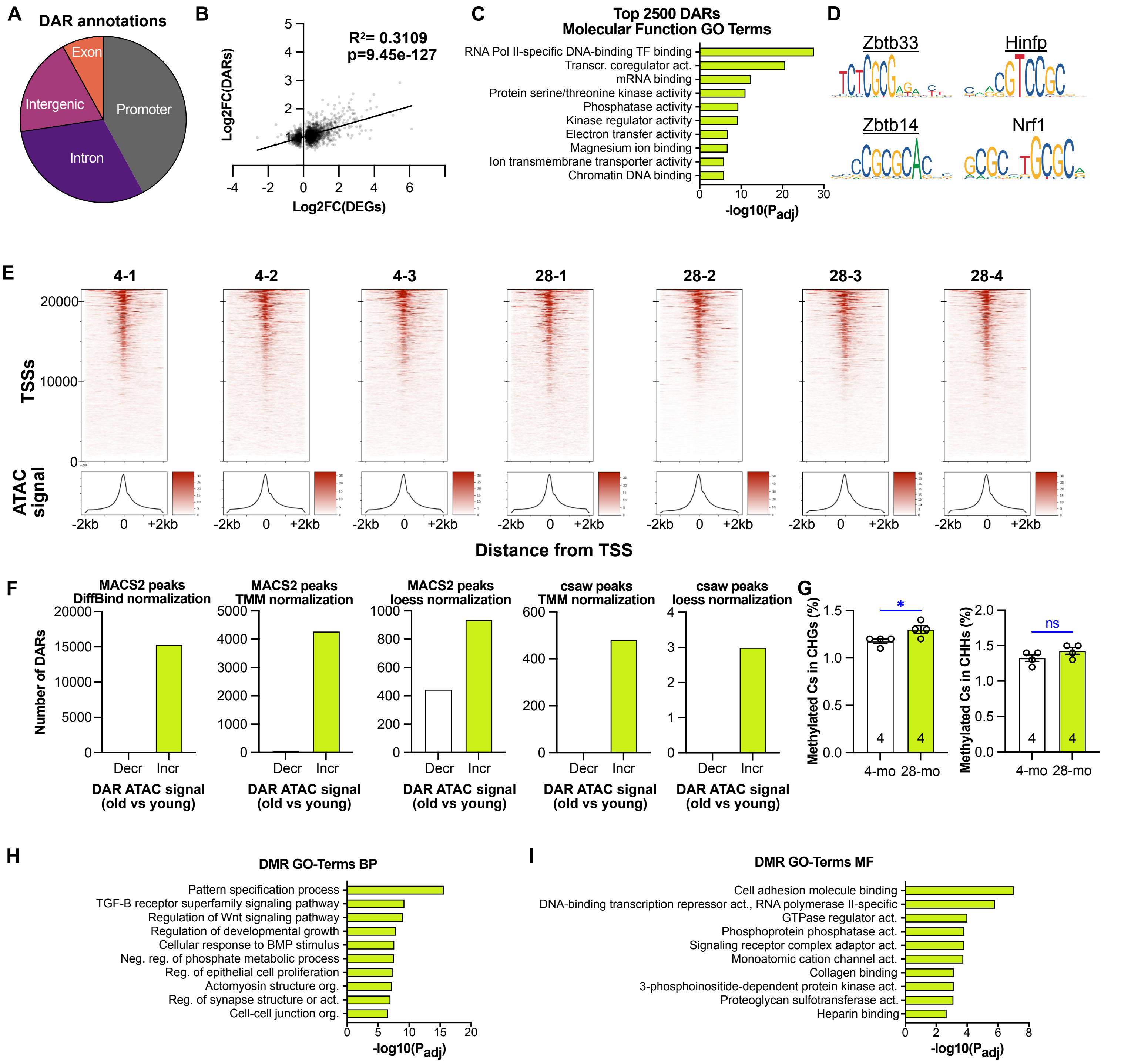


**Supplementary Figure 8. Quality control and additional analyses of chromatin accessibility data.** **(A)** Genomic annotation of snATACseq DARs, showing the distribution of accessible regions across different genomic features. **(B)** Correlation analysis between log2 fold changes of differentially accessible regions (DARs) and their associated differentially expressed genes (DEGs) from snATACseq and snRNAseq, respectively. P-value is based on the Pearson correlation. **(C)** Gene Ontology analysis of molecular functions for the top 2,500 snATACseq DARs. **(D)** Motif analysis of snATACseq DARs, highlighting four transcription factors with binding motifs enriched in CpG-rich regions. Representative motif logos are shown. **(E)** TSS enrichment analysis of bulk ATAC-seq samples from young and aged hearts. The plot shows the average ATAC-seq signal intensity around transcription start sites (TSS) for all samples, demonstrating the quality and consistency of the ATAC-seq data across different age groups. **(F)** Comparison of bulk ATAC-seq results using different normalization methods, demonstrating robustness of increased accessibility of aged chromatin regardless of the analytical method. **(G)** Percentage of methylated cytosines in CHG (left) or CHH (right) contexts in young and aged CM nuclei. H = A, T, or C. *t*-test. **(H-I)** Gene ontology biological process (BP) and molecular function (MF) terms associated with DMRs between young and aged CMs. *, P<0.05.

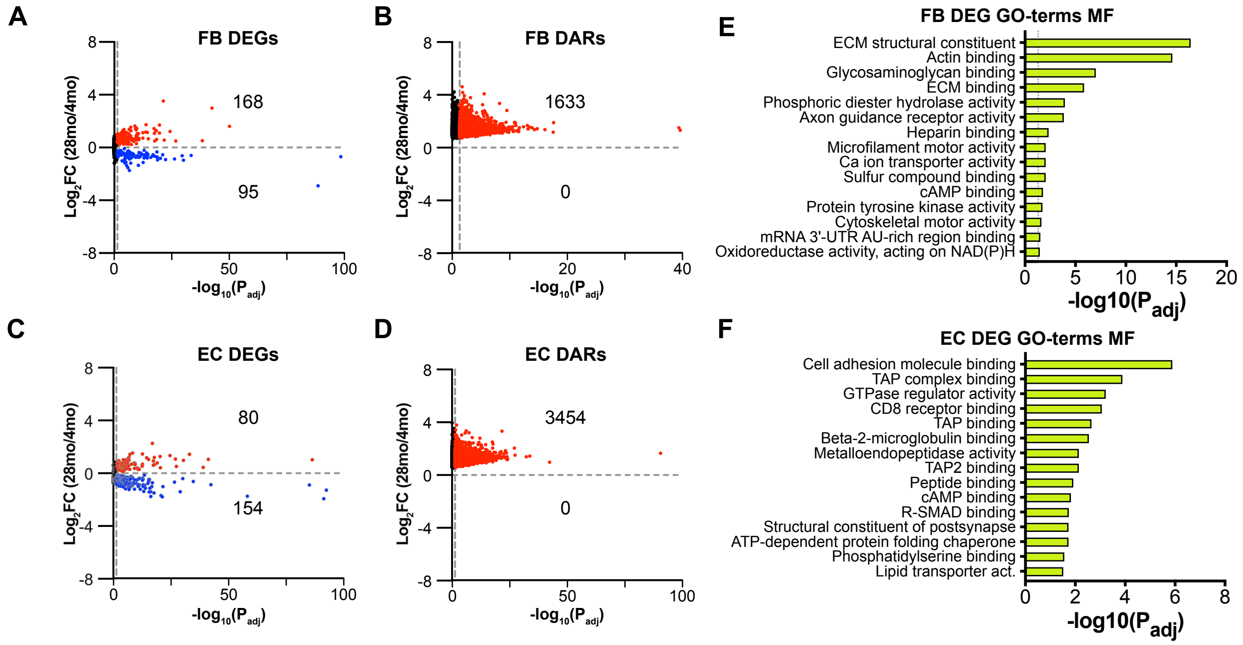


**Supplementary Figure 9. Transcriptomic and epigenomic changes in cardiac fibroblasts and endothelial cells during aging. (A-D)** Volcano plots showing DEGs (A, C) and DARs (B, D) in cardiac fibroblasts (FBs: A-B) and endothelial cells (ECs: C-D) between young and aged hearts. DEGs or DARs that were significantly up- or down- regulated are shown in red and blue, respectively. **(E-F)** Gene Ontology analysis of molecular functions (MF) associated with DEGs in FBs (E) or ECs (F).

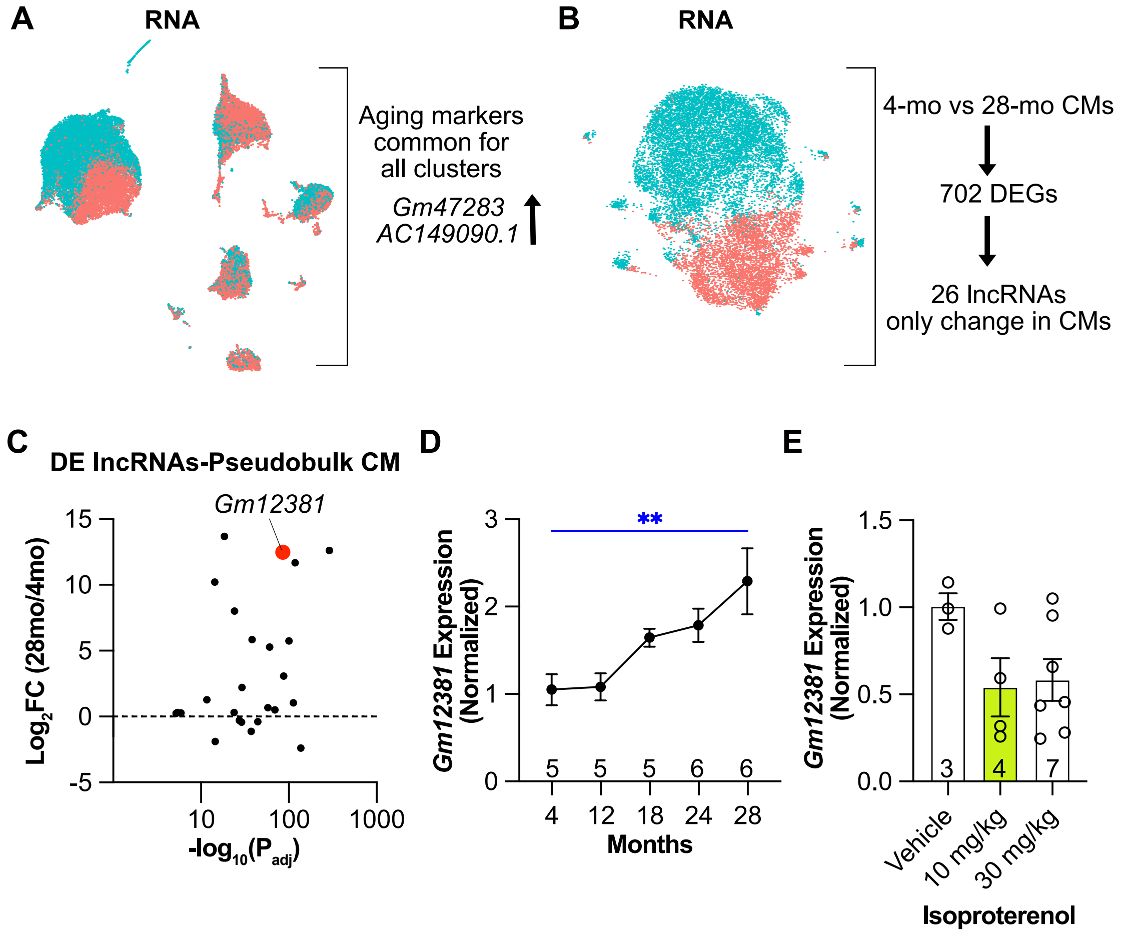


**Supplementary Figure 10. Long non-coding RNAs as potential biomarkers of cardiac aging. (A)** Schematic representation of UMAP plot highlighting the upregulation of *Gm47283* and *AC149090.1* in all aged cardiac cell types. **(B)** Schematic representation showing 26 lncRNAs that are differentially expressed exclusively in aged vs. young CMs. **(C)** LncRNA expression changes in aged CMs, highlighting *Gm12381* as one of the most significantly and robustly upregulated lncRNAs. **(D)** RTqPCR analysis of *Gm12381* expression in whole heart samples across different ages. Expression normalized to *Ppia* continuously increased with age. One way ANOVA with Dunnett's post hoc test. **(E)** *Gm12381* expression levels in hearts after treatment with vehicle (saline) or isoproterenol. *Gm12381* normalized to *Ppia* did not change in response to isoproterenol stress for 4 weeks. One-way ANOVA with Dunnett's test. Data are presented as mean ± SEM. **$p < 0.01$.

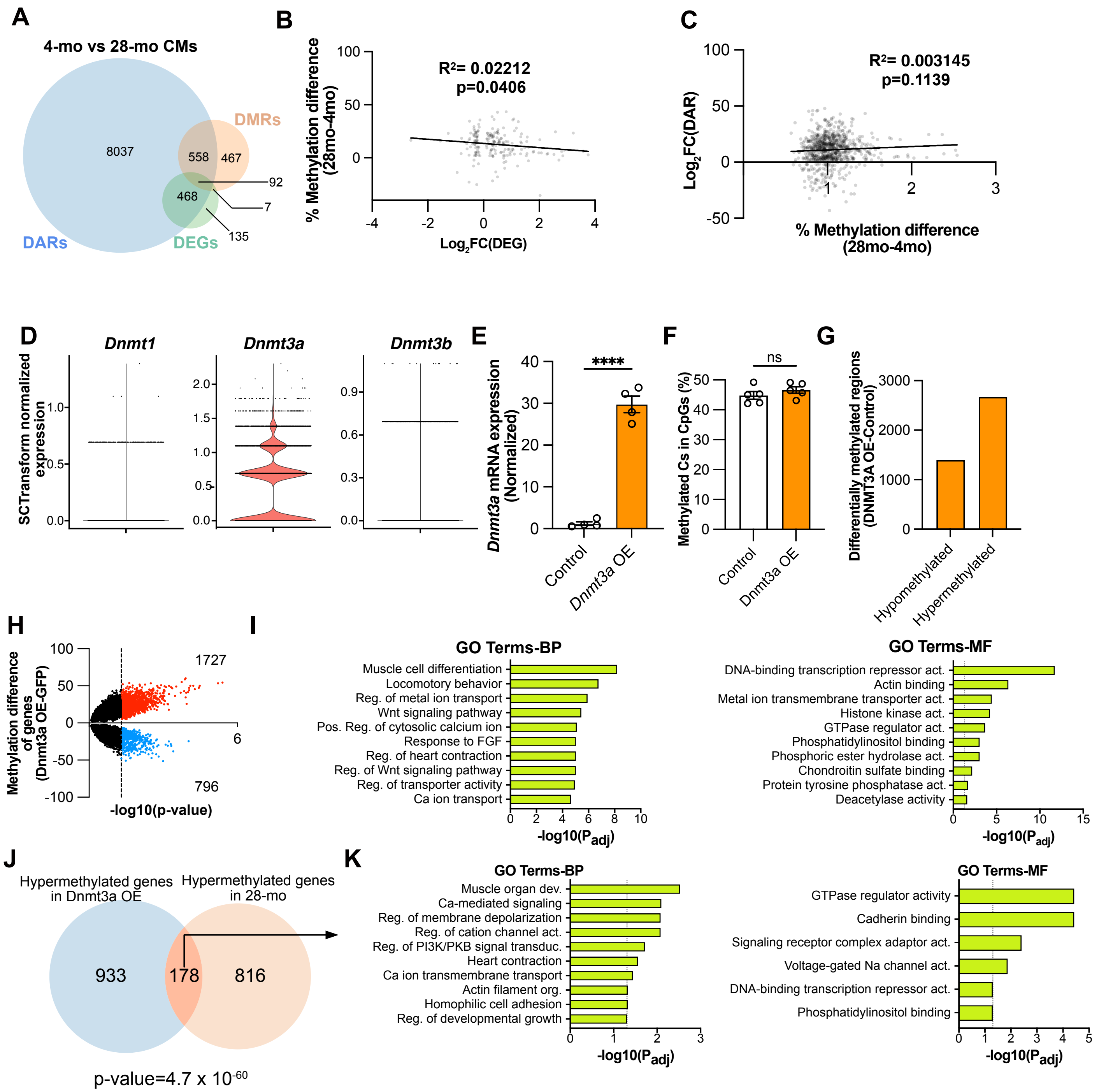


**Supplementary Figure 11. Integration of multi-omics data and *Dnmt3a* overexpression analysis.** **(A)** Venn diagram showing the overlap between DEGs and genes neighboring DARs or DMRs in aged vs. young CMs. **(B-C)** Scatter plots showing the relationship between DMRs and DARs (B) or DEGs (C). DMRs were associated with DARs when they shared the same neighboring gene. There was no significant correlation (Pearson correlation) between differential methylation and accessibility or expression. **(D)** Expression levels of different DNA methyltransferases (Dnmts) in CMs. *Dnmt3a* was the predominantly expressed isoform. **(E)** RTqPCR analysis confirming *Dnmt3a* overexpression in AAV9-Dnmt3a injected hearts compared to controls (n=4 per group). t-test. Data are presented as mean ± SEM. ****, $p < 0.0001$. **(F)** Methylated Cs in CpG islands in GFP (Control) and *Dnmt3a* OE CMs. t-test. Data are presented as mean ± SEM. ns, $p > 0.05$. **(G)** Hyper and hypomethylated regions in GFP and *Dnmt3a* OE CMs shows increased methylation in *Dnmt3a* OE CMs. **(H)** Volcano plot showing differentially methylated genes in *Dnmt3a* OE CMs. **(I)** GO-Term analysis revealed changes in pathways related to transcription and calcium transport in *Dnmt3a* OE CMs, similar to changes observed in RNA-seq of *Dnmt3a* OE CMs (Figure 4M-N). **(J)** 262 genes were common in the DMR datasets for both *Dnmt3a* vs Control and 28-mo vs 4-mo comparisons. Hypergeometric test showed significant correlation between these genes. Hypergeometric test, p value<0.001 **(K)** GO-Term analysis of the common DMRs show biological pathways related to calcium and ion transport as well as molecular pathways related to DNA binding.

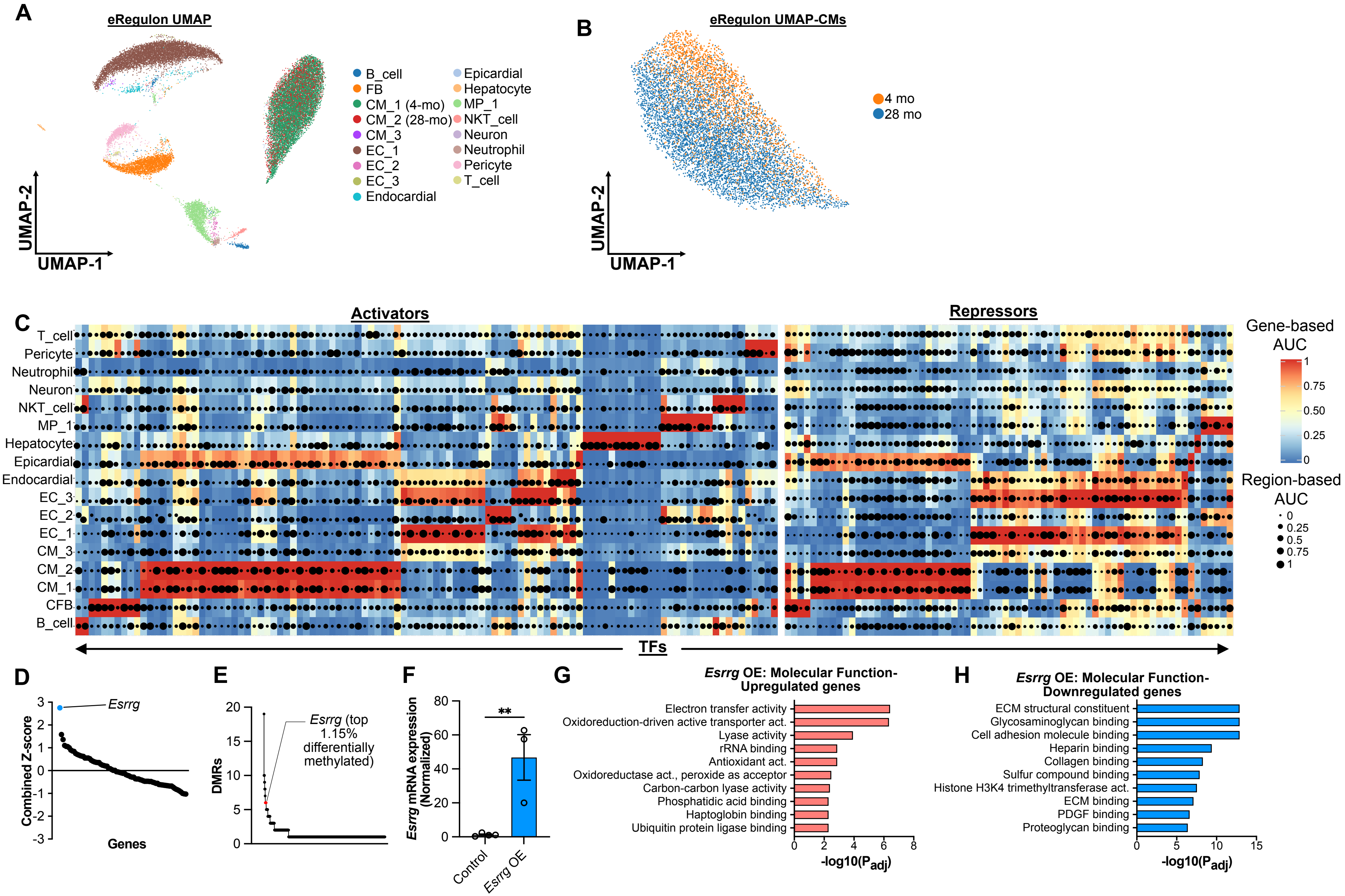


**Supplementary Figure 12. Gene regulatory network analysis and *Esrrg* methylation and overexpression.** UMAP plot showing cardiac cell population **(A)** and CM **(B)** clustering based on eRegulon profiles (right). **(C)** Heatmap displaying altered eRegulon activity between different cell types. **(D)** Z-score analysis of putative TFs that regulate gene regulatory networks of CM aging. **(E)** Histogram showing the distribution of genes based on the number of hyper methylated regions (DMRs) they contain. *Esrrg*, with 6 different methylated regions, is highlighted as being in the top 1.15% of genes with the most DMRs. **(F)** RTqPCR analysis confirming *Esrrg* overexpression in AAV9-Esrrg injected hearts compared to controls. *t*-test. Data are presented as mean ± SEM. $^{**}p < 0.01$. **(G-H)** Gene ontology analysis of molecular function for upregulated (E) and downregulated (F) DEGs.

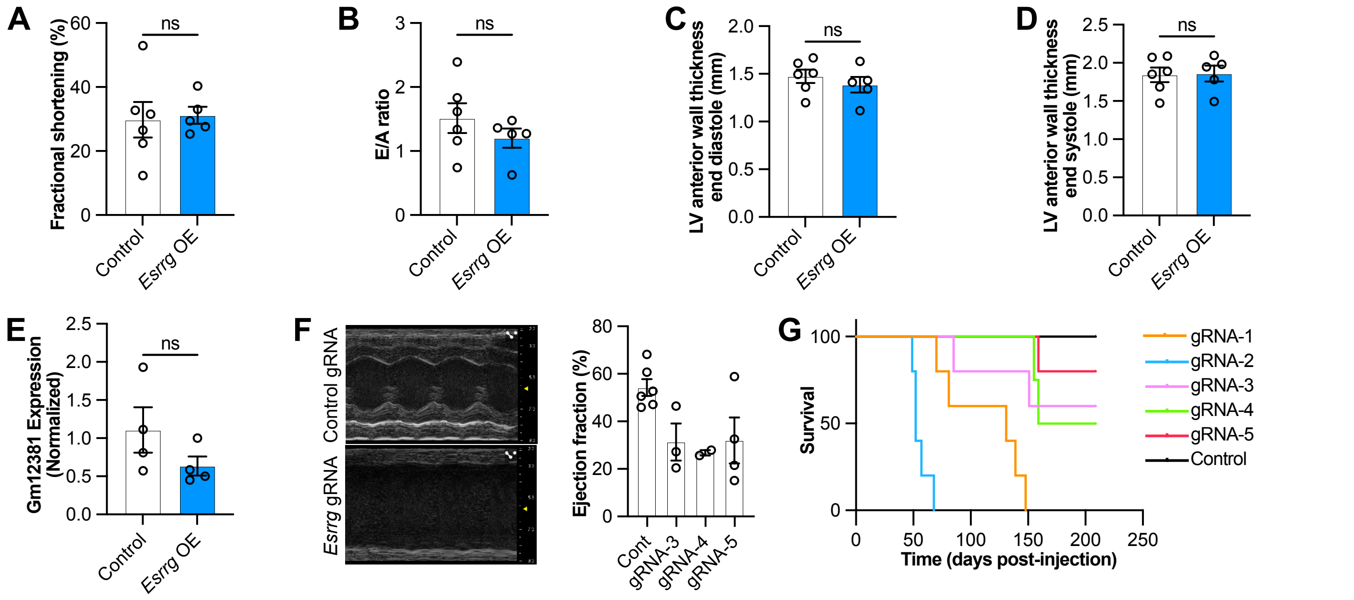


**Supplementary Figure 13. Cardiac effects of *Esrrg* overexpression in aged hearts and *Esrrg* knockout in young hearts. (A-D)** Echocardiographic assessment of 28-month-old mice 4 months after injection with MyoAAV-Tnnt2-GFP (control) or MyoAAV-Tnnt2-Esrrg (*Esrrg* OE) vectors. Fractional shortening, reflecting systolic function, E/A ratio, an indicator of diastolic function, and LV wall thickness did not significantly differ between groups. *t*-test. **(E)** Expression of aging marker *Gm12381* in control and *Esrrg* OE hearts 4-months after injection. *t*-test. **(F)** *Esrrg* deletion resulted in severe systolic dysfunction in adult mice. **(G)** Survival curve demonstrating increased mortality in *Esrrg* knockout mice compared to controls. Data are presented as mean ± SEM; ns=not significant.
